# Dissecting the common and compartment-specific features of COVID-19 severity in the lung and periphery with single-cell resolution

**DOI:** 10.1101/2020.06.15.147470

**Authors:** Kalon J. Overholt, Jonathan R. Krog, Bryan D. Bryson

## Abstract

As the global COVID-19 pandemic continues to escalate, no effective treatment has yet been developed for the severe respiratory complications of this disease. This may be due in large part to the unclear immunopathological basis for the development of immune dysregulation and acute respiratory distress syndrome (ARDS) in severe and critical patients. Specifically, it remains unknown whether the immunological features of the disease that have been identified so far are compartment-specific responses or general features of COVID-19. Additionally, readily detectable biological markers correlated with strata of disease severity that could be used to triage patients and inform treatment options have not yet been identified. Here, we leveraged publicly available single-cell RNA sequencing data to elucidate the common and compartment-specific immunological features of clinically severe COVID-19. We identified a number of transcriptional programs that are altered across the spectrum of disease severity, few of which are common between the lung and peripheral immune environments. In the lung, comparing severe and moderate patients revealed severity-specific responses of enhanced interferon, A20/IκB, IL-2, and IL-6 pathway signatures along with broad signaling activity of *IFNG, SPP1, CCL3, CCL8*, and *IL18* across cell types. These signatures contrasted with features unique to ARDS observed in the blood compartment, which included depletion of interferon and A20/IκB signatures and a lack of IL-6 response. The cell surface marker *S1PR1* was strongly upregulated in patients diagnosed with ARDS compared to non-ARDS patients in γδ T cells of the blood compartment, and we nominate S1PR1 as a potential marker for immunophenotyping ARDS in COVID-19 patients using flow cytometry.

**HIGHLIGHTS:**

- COVID-19 disease severity is associated with a number of compositional shifts in the cellular makeup of the blood and lung environments.
- Transcriptional data suggest differentially expressed cell surface proteins as markers for COVID-19 immunophenotyping from BALF and PBMC samples.
- Severity-specific features COVID-19 manifest at the pathway level, suggesting distinct changes to epithelia and differences between local and systemic immune dynamics.
- Immune-epithelial cellular communication analysis identifies ligands implicated in transcriptional regulation of proto-oncogenes in the lung epithelia of severe COVID-19 patients.
- Network analysis suggests broadly-acting dysregulatory ligands in the pulmonary microenvironment as candidate therapeutic targets for the treatment of severe COVID-19.

## INTRODUCTION

Within months of the identification of the novel coronavirus SARS-CoV-2 in Wuhan, China in December 2019, the virus had spread to every major country on Earth [1, 2, 3, 4]. The pandemic disease caused by SARS-CoV-2, termed coronavirus disease 2019 (COVID-19), has diverse clinical presentations ranging from asymptomatic infection, to moderate symptomatic infection with possible pneumonia, to severe respiratory distress, to critical respiratory failure, septic shock, and/or multiple organ dysfunction or failure [1, 4, 5, 6]. A hallmark of severe and critical COVID-19 cases is a rampant dysregulation of the immune system concomitant with the development of a hypoxemic respiratory condition widely characterized as acute respiratory distress syndrome (ARDS) [1, 7, 8, 9, 10]. The serological profile of severe COVID-19 patients largely resembles the cytokine profile of ARDS [9] and has been characterized by high levels of many cytokines including IL-2, IL-6, IL-7, IL-8, IL-18, CCL2, CCL8, TNF-α, and IFN-γ [9,11, 12, 13, 14]. Post-mortem examinations of COVID-19 patients reveal the aftermath of these disease dynamics: diffuse alveolar damage, multi-organ infiltration of lymphocytes and alveolar macrophages [1, 7, 15, 16], and pneumocyte hyperplasia and peribronchiolar metaplasia in the epithelium resembling adenocarcinomas [15, 16, 17]. These morphological findings suggest local and systemic activation and infiltration of inflammatory immune cells that may contribute to the inflammatory injuries of the respiratory system consistent with ARDS phenotypes in severe COVID-19 [15].

While months of clinical observations in hospitals across the world have led to consistent descriptions of COVID-19 severity at a clinical level [18, 19, 20], the biological underpinnings of immune hyperactivation in severe and critical COVID-19 are only beginning to be defined. Bulk RNA sequencing (bulk RNA-seq) and single-cell RNA sequencing (scRNA-seq) studies have identified stark transcriptional differences between bronchoalveolar lavage fluid (BALF) and peripheral blood mononuclear cell (PBMC) samples in hospitalized COVID-19 patients, indicating that immunological responses may be highly compartment-specific [21, 22]. A number of recent studies have compared severe patients to healthy control subjects in an effort to define immunological hallmarks of disease severity in both the lung and peripheral circulation using scRNA-seq. These studies have identified that severe disease in the lung compartment is associated with lymphopenia, T cell hyperactivation, and inflammatory macrophage polarization, while severe disease in the blood is associated with lymphopenia, suppression of type I and type II interferon activity, and decreased monocyte HLA class II expression [23, 24, 25, 26]. Although these studies describe immunological dysregulation in severe COVID-19 patients requiring intensive care, they do not explore immunological features distinguishing COVID-19 patients who experience moderate or non-ARDS pathology from those who progress to life-threatening disease courses. Single-cell resolution studies specifically comparing disease severity strata are needed to reveal the immunological mechanisms responsible for severity-specific immune dysregulation and to identify signatures of disease severity that may inform COVID-19 patient triage and treatment.

While many studies have endeavored to apply transcriptional profiling to understand the cellular dynamics underlying coronavirus infection, to our knowledge, a single integrative study comparing responses across the spectrum of disease severity in both the lung and blood compartments has not yet been reported. Isolated analyses of BALF samples have shown neutrophil and proliferating T cell enrichment in severe over moderate patients accompanied by CD8+ T cell lymphopenia, a shift towards inflammatory macrophage polarization, and increased transcriptional expression of a variety of cytokines [27, 28]. Previous investigations focusing only on peripheral blood responses have shown various compositional shifts between severe and moderate disease including cytopenias of monocytes, natural killer (NK) cells, dendritic cells (DCs), and T cells, [28, 29, 30, 31, 32] accompanied by increases in plasmablasts, B cells, and neutrophils [33, 34, 35, 36]. Serum cytokine levels are reportedly altered between severe and moderate patients, including increased IL-6, IL-8, and TNF-ɑ [11, 35] and deficient IFN-α and IFN-γ [33], while blood single-cell transcriptional analyses have shown alteration of the TNF-α/NF-κB pathway, interferon signatures, and HLA class II expression [33, 34, 36, 32].

While individual studies have increased our understanding of severe COVID-19 in the lung and blood separately, transcriptional dynamics have not been leveraged to propose detectable cell surface markers correlated with the level of a patient’s disease severity in cell types of either compartment. Another large gap in the literature persists regarding the transcriptional similarities and differences between the local and peripheral immune environments. Critically, dysregulated biological pathways and their correlation with clinical severity levels remain poorly understood in immune cells of both compartments. Immune-epithelial signaling dynamics at the site of local infection likely play a supporting role in severe responses but have also not been rigorously described. An overarching unaddressed question is the possibility for therapeutic interventions to affect the lung and blood compartments differently; thus, generating comparative transcriptional data will be useful for determining efficacious treatment strategies for severe COVID-19.

In this study, we present comparisons between immunological signatures of COVID-19 based on clinical severity level in both the lung and blood compartments using identical methods. We re-analyzed two publicly available scRNA-seq datasets: one containing BALF cells from donors stratified by the original authors as “moderate” and “severe” COVID-19 patients as well as healthy control subjects, obtained from Liao et al. [27], and one containing PBMCs from donors stratified by the original authors as “non-ARDS” and “ARDS” patients as well as healthy control subjects, obtained from Wilk et al. [32] For both of these datasets we conducted identical pre-processing, integration and analysis in order to obtain comparable results. We first evaluated differential gene expression and pathway-level changes across the disease severity stratifications mentioned above for each cell type in the BALF and the PBMC datasets. Next, we leveraged differential expression data to identify cell surface proteins that could be promising markers for COVID-19 severity immunophenotyping. Finally, we investigated ligand-receptor interactions implicated in regulating severity-specific immunophenotypes using NicheNet network analysis [37] and we propose multiple ‘pan-ligands’ as potential key regulators specific to severe disease. Our integrative analysis adds to a growing knowledge-base and contributes to a finer understanding of the mechanisms that drive ARDS-related immune dysregulation in severe COVID-19. Our findings may guide future work informing potential interventional strategies to improve patient outcomes as the COVID-19 pandemic continues to unfold.

## RESULTS

### Single cell RNA-sequencing (scRNA-seq) reveals severity-specific compositional changes across the diversity of cells in the lung microenvironment and blood

Severe COVID-19 has been thought to result in profound immune dysregulation at both a local and systemic level. Here, we performed a re-analysis of single-cell RNA sequencing (scRNA-seq) data to identify common and compartment-specific signatures of COVID-19 disease severity. We used identical methods to separately analyze multi-donor scRNA-seq datasets from bronchoalveolar lavage fluid (BALF) and peripheral blood mononuclear cells (PBMCs) in COVID-19 patients classified by severity strata as well as healthy control subjects to investigate severity-specific immune dysregulation in the lung and periphery. BALF and PBMC raw gene-barcode matrices were downloaded from separate studies in the GEO [27, 32]. The BALF dataset consisted of 9 patients (6 severe and 3 moderate) and 4 healthy control donors, while the PBMC dataset consisted of 7 patients (4 patients with ARDS, 4 patients without ARDS, where one patient who was sampled twice fell within both groups at different stages of the disease course) and 6 healthy control donors. We performed our analyses on BALF and PBMC datasets separately with the goal of identifying common and compartment-specific gene signatures.

To analyze the compositional changes in cell populations occurring during severe COVID-19, we first sought to identify clusters in transcriptionally heterogeneous scRNA-seq data of the lung. After preprocessing raw BALF gene-barcode matrices, a filtered dataset consisting of 66,452 cells was clustered using Seurat and visualized by uniform manifold approximation and projection (UMAP), as shown in **Figure 1A**. A total of 33 clusters were identified showing distinct separation in a two-dimensional UMAP space. Annotation of these clusters based on the expression of canonical cell type markers showed the presence of myeloid cells, lymphoid cells, and epithelial cells, closely matching the cell types found by the original publication [27]. Of 13 annotated cell types, 12 contained cells from moderate, severe, and control donors, with the exception of plasma cells which were not recovered from moderate donors (**Figure S1A**). Compositional changes across disease severity categories were assessed by z-scored percentages of cell types in each donor sample. Analyzing BALF compositional changes showed trends of increased donor specific percentages of NK cells, mixed T cells, and plasmacytoid dendritic cells (pDCs) in moderate donors over control and severe donors as displayed in the **Figure 1A** heatmap.

**Figure 1.**
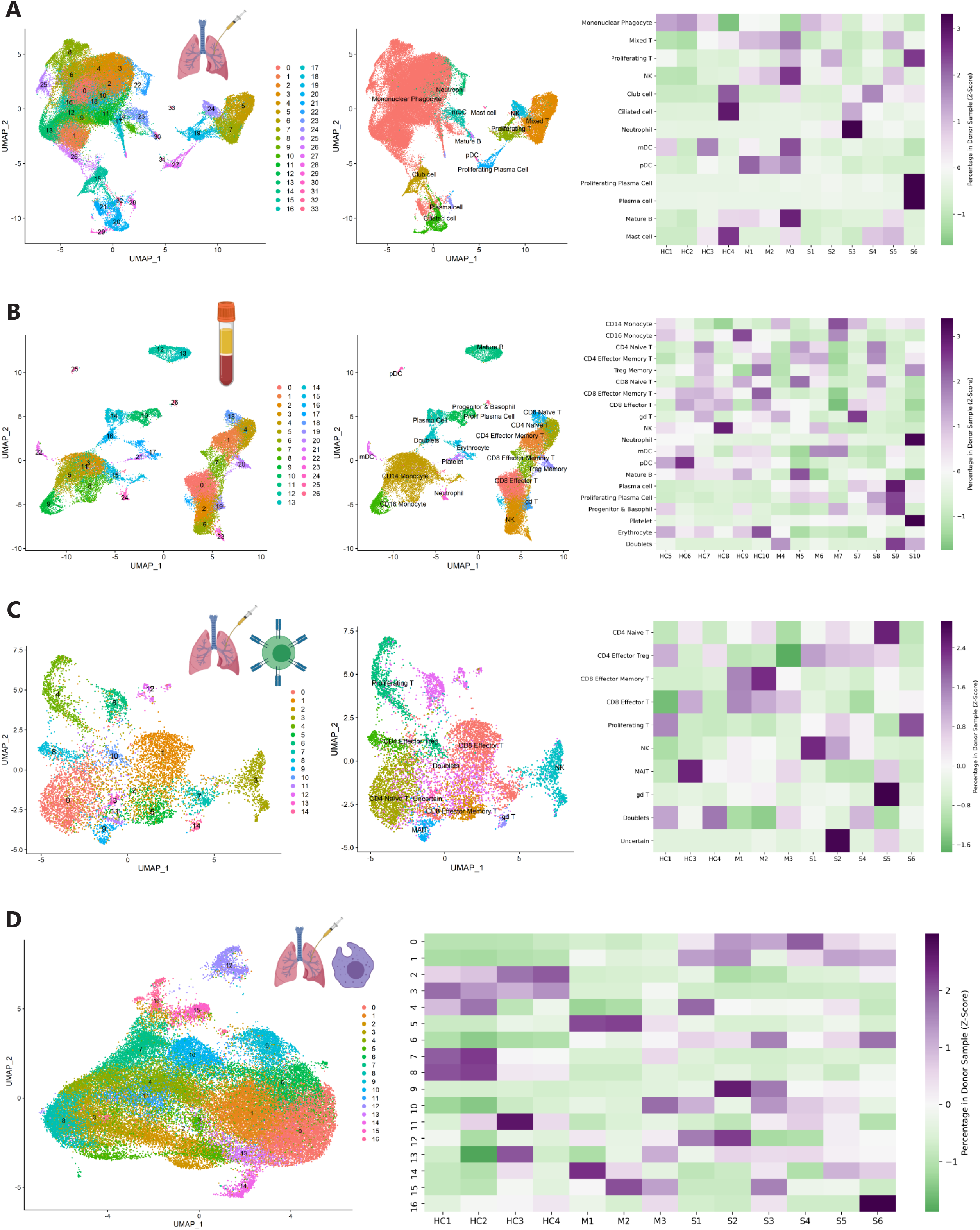
Single cell RNA sequencing reveals cellular diversity of bronchoalveolar lavage (BALF) fluid and peripheral blood mononuclear cells (PBMCs) in healthy controls and across the spectrum of COVID-19 severity. **A) (Left)** UMAP projection and clustering of 66,452 cells identified in BALF across healthy (*n*=4, HC1-HC4), moderate (*n*=3, M1-M3), severe (*n*=6, S1-S6) donors. (**Middle)** Annotation of BALF clusters following merging into 13 cell types. pDC=plasmacytoid dendritic cell, mDC=myeloid dendritic cell, NK=natural killer cell. (**Right)** Percent composition (z-score) of BALF cell types in each donor sample. **B) (Left)** UMAP projection and clustering of 31,957 cells identified in peripheral blood mononuclear cells (PBMCs) across healthy (*n*=6, HC5-10), non-ARDS (*n*=4, M4-M7), and ARDS (*n*=4, S7-S10) donors. **(Middle)** Annotation of PBMC clusters following merging into 19 cell types. Prolif Plasma Cell=proliferative plasma cell, gd T= gamma delta T cell. **(Right)** Percent composition (z-score) of PBMC cell types in each donor sample **C) (Left)** UMAP projection and clustering of BALF “NK”, “Proliferating T”, and “Mixed T” populations across healthy (*n*=3), moderate (*n*=3), and severe (*n*=5) donors following iterative data integration. **(Middle)** Annotation of BALF NK and T cell types following iterative data integration. MAIT=mucosal-associated invariant T cell. **(Right)** Percent composition (z-score) of BALF NK and T cell types in each donor sample. **D) (Left)** UMAP projection and clustering of mononuclear phagocyte cell population from the BALF across healthy (*n*=4), moderate (*n*=3), and severe (*n*=6) donors. **(Right)** Percent composition (z-score) of BALF mononuclear phagocytes in each donor sample.

We next preprocessed raw PBMC gene-barcode matrices, resulting in a filtered dataset of 31,957 cells. The blood single-cell landscape was visualized by UMAP (**Figure 1B**). A total of 26 clusters were identified, showing distinct populations in a two-dimensional UMAP space. Annotation of these clusters revealed the presence of myeloid cells and lymphoid cells, roughly matching the cell types found by in the original publication using a different method [32]. We termed the blood cells observed in this dataset as “PBMCs”, although cell type annotation revealed the presence of anuclear platelets and erythrocytes, as well as polymorphonuclear basophils and neutrophils. All 20 annotated cell types contained cells from ARDS, non-ARDS, and control donors (**Figure S1D**). Cluster 16 (*CD14*+/*CD3D*+/*IGHG4*+) was annotated as probable doublets using the marker gene panel shown in **Figure S1F**. Analyzing PBMC compositional changes across severity categories using z-scored percentages as described above, we observed trends indicating a decrease in erythrocyte and CD8+ effector memory T cell percentages as well as an expansion of plasma cells and proliferating plasma cells in both ARDS and non-ARDS patients compared to control donors.

To examine the population of NK and T cells in the BALF at finer resolution, we performed iterative data integration on the “NK”, “Mixed T”, and “Proliferating T” populations, using 5 severe donors, 3 moderate donors, and 3 healthy control donors. Severe donor S3 and healthy control donor HC2 were omitted as a consequence of having too few NK and T cells to allow integration and alignment. After re-integration, a total of 8,832 NK and T cells were visualized via UMAP (**Figure 1C**). A total of 14 clusters were identified, which were annotated according to 8 distinct cell type labels with additional cluster 13 labeled as “uncertain” and clusters 6 (*CD3D*+/*CD68*+) and 12 (*CD3D*+/*FCGR3B*+) classified as probable doublets using the marker gene panel shown in **Figure S1I**). Analyzing NK and T cell compositional changes across severity categories, we observed trends indicating expansion of CD4+ naive T and CD4+ effector Treg cells in moderate donors compared to severe and control donors.

The distinctive heterogeneity observed in mononuclear phagocytes (MPs) of the BALF seen in **Figure 1A** led us to address severity-specific changes in individual MP populations. We again performed iterative data integration on 19 clusters identified as MPs in the original BALF dataset (**Figure 1A**). Following iterative data integration using 6 severe donors, 3 moderate donors, and 4 healthy controls, a total of 50,146 MP cells were visualized by UMAP (**Figure 1D**). The 16 MP clusters were left unannotated in further analysis. The clusters were evaluated using a panel of BALF marker genes to identify probable doublets (**Figure S1L**), and clusters 12 (*CD68*+/*CD3D*+) and 16 (*CD68*+/*IGHG4*+) were classified as such. Analysis of compositional changes of MP clusters across disease categories showed more pronounced compositional changes than we observed in the BALF in general. Clusters 0, 1, and 6 (expressing chemokines *CCL2*, *CCL3*, *CCL8* as well as *SPP1*) appear to be expanded in severe donors compared to moderate and control donors, while clusters 2, 3, and 8 (expressing *MRC1*, *C1QA*, and *FABP4*) appear to be expanded in control donors compared to moderate and severe donors. Cluster 5 (expressing *HLADQA1* and *HLADQA2*) appears to be uniquely expanded in moderate donors compared to control and severe donors.

### Transcripts coding for cell surface proteins are differentially expressed across the COVID-19 severity spectrum and may be useful as markers for immunophenotyping

Cellular markers corresponding to clinical categories of COVID-19 disease severity have not yet been rigorously characterized. Here, we sought to leverage transcriptional differences across severity strata to identify surface-bound proteins in both the lung and blood that could be used as markers for immunophenotyping. To nominate putative cell surface markers, we conducted differential gene expression analysis between severe and moderate patient groups for lung (BALF) cells and between ARDS and non-ARDS patient groups for blood (PBMC) cells and selected significant differentially expressed genes (DEGs) with a fold change (FC) cutoff of |log_2_FC>1|. **Figure 2A** shows significantly up- and down-regulated transcripts for pDCs in the BALF, one of 17 lung cell types studied. From the large number of significant DEGs, we observed the upregulation of *AREG* and *CD55*, which were verified as cell surface protein-coding genes using the Cell Surface Protein Atlas [38]. The *AREG* and *CD55* transcripts show robustly increased expression in pDCs in severe compared to moderate disease (**Figure 2B**) and compared to pDCs in healthy control subjects (**Figure S2A**). We additionally showed that *AREG* and *CD55* are not upregulated in moderate disease compared to controls (**Figure S2A**). The expression of *AREG* and *CD55* was cross-referenced with bulk RNA-seq blood data from the human Immune Cell Atlas (**Figure S2C**), and expression of these transcripts in various subtypes of DCs was verified [39, 40]. Differential expression of transcripts coding for cell surface proteins was evaluated for every cell type in the BALF, resulting in a matrix of potential severity-specific surface markers for each cell type (**Figure 2C)**. Upregulation of the *AREG* gene was also observed in myeloid dendritic cells (mDCs, also known as conventional dendritic cells), indicating a possible conserved response across DC subtypes. Extending our analysis beyond immune cell types and relaxing the stringency cutoff of |log_2_FC>1|, we made the interesting observation that epithelial (club and ciliated) cells demonstrate significant upregulation of *ICAM1* and *LDLR*, the two main entry receptors for respiratory rhinoviruses. We confirmed that markers identified as differentially regulated between severe and moderate disease were not differentially regulated in the same direction between moderate disease and control (**Figure S2B**), with the exception of *HLADRA* and *VAMP5* on MPs and *CCR7* on mDCs, indicating that all other identified cell surface protein transcripts in **Figure 2C** are unique markers of severe disease.

**Figure 2.**
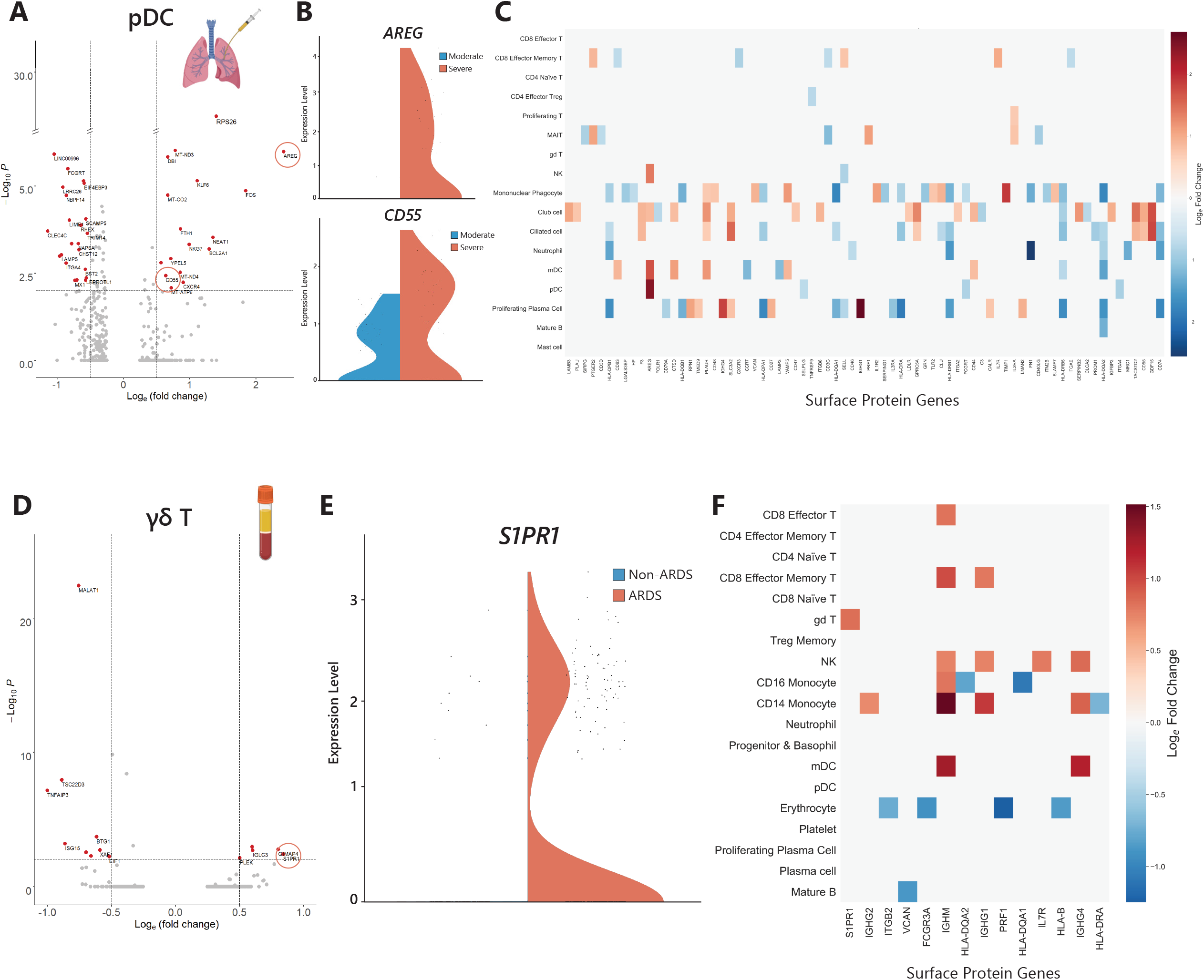
Transcriptional signatures of COVID-19 severity include surface markers in cells of the lung microenvironment and peripheral blood. **A)** Differentially expressed genes were identified in 17 of 18 cell types found in the BALF between moderate and severe COVID-19 patients. An example volcano plot of differentially expressed genes is shown for plasmacytoid dendritic cells (pDCs). Genes denoted by red dots meet a threshold of p<0.01 and natural log fold change (ratio of severe to moderate expression) greater than 0.5. The circled cell surface markers *AREG* and *CD55* are significantly unregulated in severe disease in pDCs. **B)** Transcriptional expression levels of the cell surface markers *AREG* and *CD55* in pDCs of moderate COVID-19 patients (blue violins, *n*=3) and severe COVID-19 patients (red violins, *n*=6). **C)** Differentially expressed surface markers for all cell types identified in the BALF (moderate donors: *n*=3, severe donors: *n*=5 for T and NK cells, *n*=6 for all other cell types). Only genes with significant differential expression (|log_2_FC|>1 and adjusted p<0.05) are shown. **D)** Differentially expressed genes were identified for all 19 of the cell types found in PBMCs between non-ARDS and ARDS-diagnosed COVID-19 patients. An example volcano plot of differentially expressed genes is shown for gamma delta (γδ) T cells. Genes denoted by red dots meet a threshold of p<0.01 and natural log fold change greater than 0.5. The circled cell surface marker *S1PR*1 is significantly upregulated in γδ T cells of ARDS patients compared to non-ARDS patients. **E)** Transcriptional expression levels of the cell surface marker *S1PR1* in non-ARDS COVID-19 patients (blue violins, *n*=4) and ARDS-diagnosed COVID-19 patients (red violins, *n*=4). **F)** Differentially expressed surface markers for all PBMC cell types identified in the peripheral blood (non-ARDS donors: *n*=4, ARDS donors: *n*=4). Only genes with significant differential expression (|log_2_FC|>1 and adjusted p<0.05) are shown.

We also sought to identify ARDS-specific cell surface markers in the blood in order to nominate immunophenotyping markers on cells that may be accessible through a blood draw. In gamma delta T (γδ T) cells, we observed ARDS-specific differentially expressed genes including the surface marker *S1PR1* (**Figure 2D**), a G-protein coupled receptor that interacts with multiple inflammatory pathways such as JAK/STAT and mTOR/PI3K/Akt [41, 42]. *S1PR1* expression was robustly upregulated in ARDS compared to non-ARDS patients (**Figure 2E**) and ARDS compared to control (**Figure S2D**), and additionally did not show upregulation between non-ARDS patients and controls. Similarly to our analysis of the BALF, we analyzed differential expression of surface protein genes in every cell type of the PBMC pool, and the matrix of potential markers is shown in **Figure 2E**. We found that *S1PR1* was also differentially expressed in CD8+ effector T cells, and its general expression on T cells was verified in bulk RNA-seq data (**Figure S2F**). Immunoglobulin heavy chain genes also appeared to be differentially expressed in unexpected blood cell types. In both BALF and PBMCs, we observed that increasing disease severity was associated with the common downregulation of HLA class II genes *HLADQA1* and *HLADQA2* on CD16+ monocytes and BALF MPs and *HLADRA* on CD14+ monocytes and BALF MPs. Overall, these results suggest cell surface proteins whose transcripts are differentially expressed across disease severity categories as candidate immunophenotyping markers in cells of the lung and blood.

To complement our transcriptomic analysis, we employed gene ontology (GO) analysis to identify the biological functions of DEGs (**Figure S2G-I**). GO analysis of the DEGs in BALF and PBMCs shows differential expression of transcripts belonging to ontologies of the innate immune system, cytokine signaling, and adaptive immune cell proliferation and activation.

### Pathway-level regulation differentiates COVID-19 severity categories, with distinct differences arising between responses in the pulmonary microenvironment and systemic circulation

To probe the molecular mechanisms of severe COVID-19 at a broader scale, we sought to utilize the large number of differentially expressed transcripts to identify biological pathways that are altered across the spectrum of disease severity in the local lung microenvironment (BALF) and systemic circulation (PBMCs). Significant transcriptional changes in cells of the BALF were abundant in all cell types with the exception of plasma cells. We first directed our attention to the epithelium, where substantial morphological damage and histological atypia were observed in deceased COVID-19 patients [1, 7, 15, 17, 16]. At the level of individual gene regulation, we first noted a surprisingly strong significant upregulation of *FOS* and *JUN*, which code for proteins implicated in the development of cancer (**Figure 3A**). Additionally, we noted a strong significant downregulation of the tumor suppressor gene *C2orf40*. When comparing ciliated cells between moderate disease and healthy control subjects (**Figure S3A**), we did not observe differential expression of proto-oncogenic transcripts. We next leveraged gene set enrichment analysis (GSEA) to dissect pathway-level regulation. GSEA provides positive and negative enrichment readouts; here, we define pathways with positive normalized enrichment score as “enriched” and those with a negative normalized enrichment score as “depleted”. GSEA revealed the enrichment of pathways involved in the innate immune response, general inflammatory response, and, surprisingly, oncogenic signaling in severe versus moderate donors. Specifically, we observed the significant enrichment of the epithelial-to-mesenchymal transition, K-Ras, PI3K/Akt and p53 pathways in addition to the TNF-α signaling via NF-κB and IL-2/STAT5 pathways.

**Figure 3.**
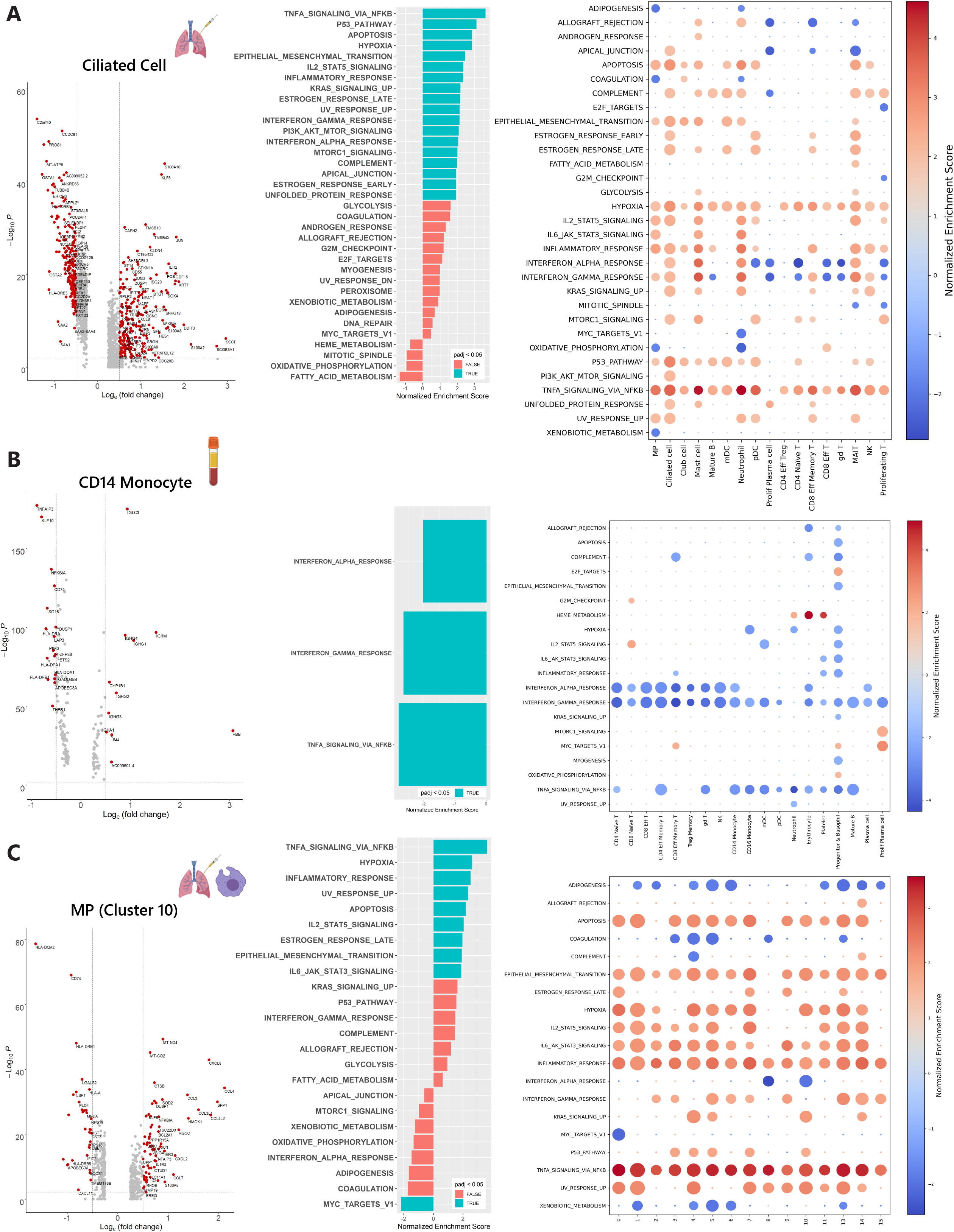
Transcriptional changes across COVID-19 severity levels indicate dysregulation of molecular pathways in cells of the blood and lung microenvironment. **A) (Left)** Volcano plot showing significantly differentially expressed genes (red dots) in ciliated cells from BALF when comparing severe and moderate patients. **(Middle)** Gene set enrichment analysis (GSEA) of hallmark gene sets in the ciliated cells of the lung. Bars colored blue are significantly enriched or depleted (adjusted p-value <0.05) while bars colored red do not pass the significance threshold. **(Right)** Hallmark gene sets indicate severity-specific enrichment of molecular pathways for all cell types found in the BALF. Normalized enrichment score is shown by a blue-red color scale and dot size is proportional to −log_10_(p), with the smallest dot size indicating non-significant adjusted p-values (p<0.05). MP=mononuclear phagocytes. **B) (Left)** Volcano plot showing significantly differentially expressed genes (red dots) in CD14+ monocytes from blood when comparing non-ARDS and ARDS-diagnosed patients. **(Middle)** Gene set enrichment analysis (GSEA) of hallmark gene sets in CD14+ monocytes. Bars colored blue are significantly enriched or depleted (adjusted p-value <0.05) in ARDS while bars colored red do not pass the significance threshold. **(Right)** Hallmark gene sets indicate ARDS-specific enrichment of molecular pathways for all PBMC cell types found in the blood. **C) (Left)** Volcano plot showing significantly differentially expressed genes (red dots) in the one of 16 mononuclear phagocyte populations (cluster 10) from the BALF when comparing severe and moderate patients. **(Middle)** Gene set enrichment analysis (GSEA) of hallmark gene sets in cluster 10 of BALF mononuclear phagocytes. Bars colored blue are significantly enriched or depleted (adjusted p-value <0.05) while bars colored red do not pass the significance threshold. **(Right)** Hallmark gene sets indicate severity-specific enrichment of molecular pathways for all mononuclear phagocyte clusters found in the BALF. Clusters 12 and 16 represent suspected doublets.

When we increased this analysis to include all of the cell types found in the BALF (**Figure 3A**), we found enriched pathways including hypoxia, TNF-α signaling via NF-κB (involving *TNFAIP3* and *NFKBIA*), and general inflammatory response to be conserved across many BALF cell types. The IFN-α and IFN-γ pathways showed a mixed response among cell types, including enrichment of these pathways in ciliated cells, mast cells, mucosal-associated invariant T cells (MAITs), and neutrophils, but depletion in proliferating plasma cells and most T cell populations Additionally, IL-2/STAT5 signaling was enriched in MPs, ciliated cells, mast cells, neutrophils, pDCs, MAITs and proliferating T cells, and IL-6/JAK/STAT3 was enriched in MPs, mast cells and neutrophils. When comparing the BALF pathway-level response between moderate disease and healthy controls (**Figure S3A**), we observed enrichment of the TNF-α signaling via NF-κB pathway in MPs, mature B cells, mDCs, neutrophils, pDCs, CD4+ effector Tregs, and NK cells. The IFN-α and IFN-γ pathways showed strong enrichment in all BALF cell types. IL-2/STAT5 signaling was enriched in MPs, club cells, mature B cells, mDCs, neutrophils, pDCs, and CD8+ effector memory T cells, while IL-6/JAK/STAT3 signaling was enriched in MPs, mature B cells, mDCs, neutrophils, and pDCs. When comparing the BALF pathway-level response between severe disease and healthy control donors (**Figure S3D**), we observed enrichment of the IFN-α and IFN-γ pathways across every cell type in the BALF.

We next investigated pathway-level changes occurring in PBMCs and found that differential gene expression between ARDS and non-ARDS patients supported the detection of statistically enriched pathways through GSEA. We first examined CD14+ blood monocytes, a cell type whose severe disease response could be compared to that of BALF MPs. In ARDS patients, CD14+ monocytes demonstrated pronounced downregulation of *ISG15*, *TNFAIP3*, and *NFKBIA* compared to non-ARDS patients, manifesting in the depletion of the IFN-α, IFN-γ, and TNF-α signaling via NF-κB pathways (**Figure 3B**). Expression of the immunoglobulin G and M heavy chains appeared to be altered at the transcriptional level but did not contribute to significantly enriched pathways. When expanding this analysis to all PBMC cell types (**Figure 3B**), we observed a striking depletion of IFN-α, IFN-γ, and TNF-α signaling via NF-κB pathways in a conserved trend across nearly every cell type. The IL-2/STAT5 and IL-6/JAK/STAT3 pathways did not appear to be significantly altered in the majority of cell types. Comparing PBMCs of non-ARDS donors to healthy controls revealed a strong enrichment of the IFN-α, IFN-γ, and TNF-α signaling via NF-κB pathways, as well as some inclusion of the IL-2/STAT5 and IL-6/JAK/STAT3 pathways (**Figure S3B**). Interestingly, comparing ARDS donors to healthy controls showed that these same pathways exhibited weaker enrichment (**Figure S3E**).

To establish a comparison between blood monocytes and BALF MPs and explore the heterogeneity of BALF MPs observed in our dataset, we analyzed differential expression and pathway-level responses across 14 of the 16 clusters resulting from iterative data integration of the MP population. Two clusters containing probable doublets were excluded from the analysis, namely cluster 12 (*CD68*+/*CD3D*+) and cluster 16 (*CD68*+/*IGHG4*+). Differential gene expression in MP cluster 10 is shown as a representative example result (**Figure 3C)**; this population exhibits upregulation of *CXCL8*, *CCL3*, *CCL4*, and *SPP1*, as well as *NFKBIA* and *TNFAIP3*. Pathway-level analysis of cluster 10 showed enrichment of TNF-α signaling via NF-κB, IL-2/STAT5 signaling, IL-6/JAK/STAT3 signaling, among others. Examining the transcriptionally heterogeneous subpopulations of MPs revealed a surprisingly homogeneous response at the pathway level. In particular, apoptosis, epithelial-to-mesenchymal transition, hypoxia, IL-2/STAT5, IL-6/JAK/STAT3, and TNF-α signaling via NF-κB were significantly enriched in severe compared to moderate disease in all or many of the clusters. The IFN-γ response appeared to be variable among clusters but trending toward enrichment. Comparison of MPs in moderate donors to healthy controls showed that most of the same pathways are broadly enriched, with the exception of epithelial-to-mesenchymal transition and hypoxia (**Figure S3C**). Further comparison between severe donors and healthy controls demonstrated a homogenous enrichment of many pathways related to cytokine signaling and innate immune responses (**Figure S3F**).

To nominate potential transcription factors related to differentially expressed gene sets, we sought to identify transcription factors linked to DEG lists using the ENCODE and ChEA ChIP-X databases. Consistent with the identified activity of the IL-6/JAK/STAT3 pathway, we observed enrichment of *STAT3* in many cells of the BALF (**Figure S3G**). *CEBPB* and *CEBPD* appeared to be depleted in the PBMC but enriched in the BALF, and opposite directionality of these genes was observed between MPs and CD14+ monocytes in particular. *NELFE* appeared to be enriched in T cells of the BALF and PBMC (**Figure S3G-H**), representing a commonality between the lung and blood responses to increasing disease severity.

### Broadly-acting dysregulatory cues induce severity-specific transcriptional changes in the lung microenvironment through receptor-ligand interactions

Following the identification of severity-specific pathway-level regulation differentiating severe and moderate disease courses, we sought to construct a putative network for how these transcriptional programs could be induced by soluble and surface-level cell-cell interactions. Specifically, we aimed to identify ligands acting as potential key regulators of severe disease in many cell types of the lung in order to nominate targets for further study as therapeutic options for severe COVID-19. **Figure 4A** shows a flow diagram detailing the steps carried out to nominate severity-specific “pan-ligands” local to the site of viral infection. Briefly, differential gene expression between moderate and severe donors was first evaluated for each cell type in the BALF. We classified these transcripts as “target genes” whose differential expression is potentially regulated through ligand-receptor interactions [37]. Next, we employed NicheNet to identify potential ligands linked to regulation of these differentially expressed target genes [37]. We applied two filtering criteria to nominate potential ligands: Ligands should (1) act on over one-third of the cell types in the BALF and (2) be differentially expressed by at least one cell type in severe disease.

**Figure 4.**
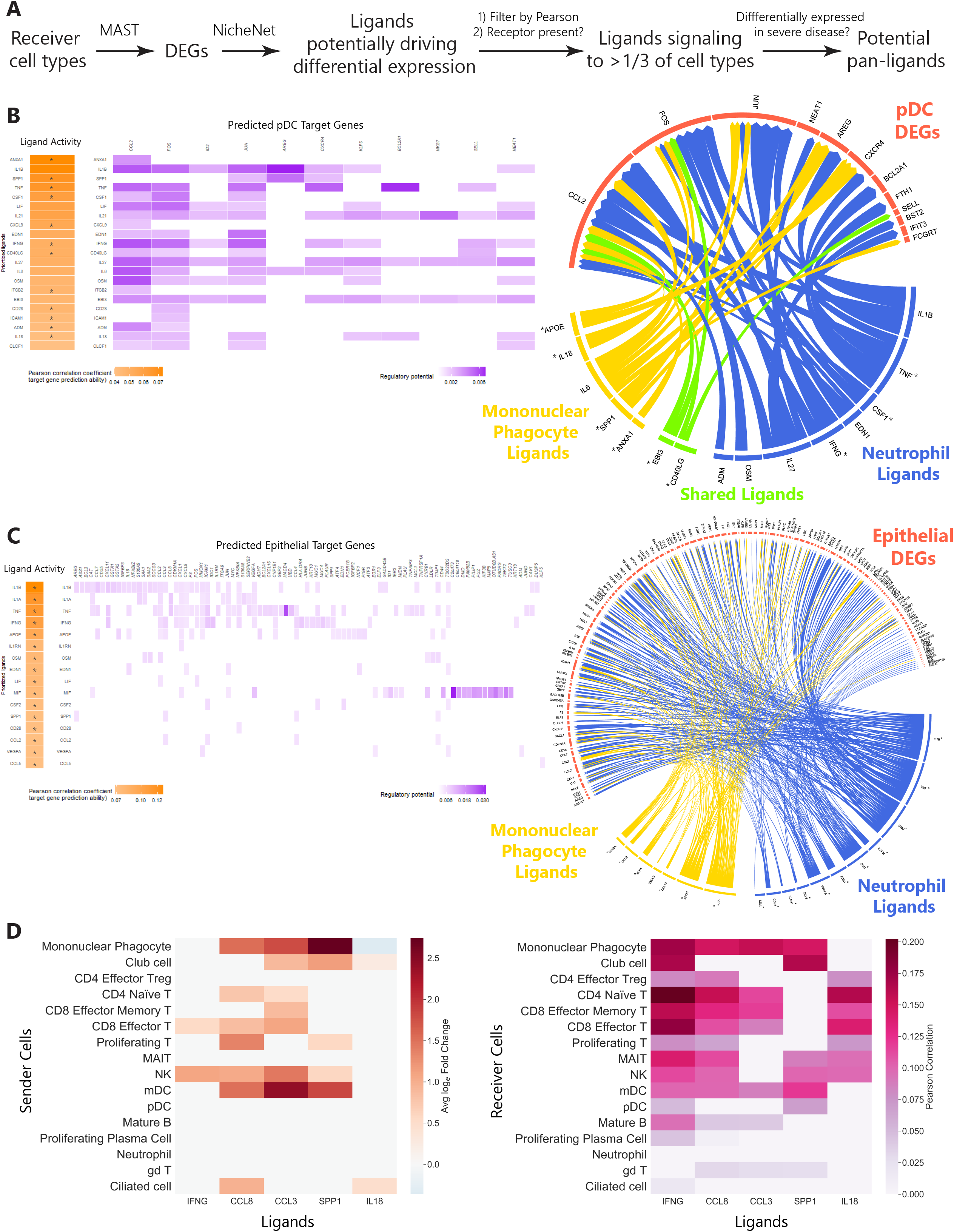
Ligand-receptor interaction networks drive severity-specific changes in gene expression in the lung microenvironment. **A)** Methods flow schematic for the identification of severity-specific pan-ligands linked to differential expression in over one-third of cell types in the BALF. **B) Left)** Predicted ligand activity driving differentially expressed genes (DEGs) in plasmacytoid dendritic cells (pDCs) in severe disease ranked by Pearson correlation coefficient (orange color scale). The potential of each ligand for inducing differential expression of pDC target genes is shown by regulatory potential (purple color scale). Asterisks (*) indicate ligands whose receptors were manually validated at the transcript level on pDCs in severe donors. **(Right)** Circos plot of ligand-target relationships between DEGs in pDCs (red) and ligands expressed by mononuclear phagocytes (yellow), neutrophils (blue), and shared ligands between both cell types (green). **C) (Left)** Predicted ligand activity driving differentially expressed genes in epithelial cells (ciliated cells and club cells) in severe disease ranked by Pearson correlation coefficient (orange color scale). The potential of each ligand for inducing differential expression of epithelial target genes is shown by regulatory potential (purple color scale). Asterisks (*) indicate ligands whose receptors were manually validated at the transcript level on epithelial cells in severe donors. **(Right)** Circos plot of ligand-target relationships between DEGs in epithelial cells (red) and ligands expressed by mononuclear phagocytes (yellow) and neutrophils (blue). **D) (Left)** “Sender” cells differentially express potential pan-ligands in the BALF between severe and moderate disease. Colored ligand-cell pairings indicate cell types in which the corresponding ligand is significantly differentially expressed in severe disease with adjusted p<0.05. Average natural log fold change (ratio of severe to moderate expression) is shown in the blue-red color scale. **(Right)** Potential pan-ligands implicated in severe COVID-19 are correlated with differential gene expression in over one-third of “receiver” cell types in the BALF. The pink color scale indicates Pearson correlation, a measure of a ligand’s ability to induce the differential gene expression measured in the corresponding “receiver” cell.

To follow up on our observation of *AREG* upregulation in pDCs during severe disease, we used NicheNet to nominate potential soluble or cell-surface mediators of pDC differential gene expression. Ligand activities were ranked using a NicheNet-generated Pearson correlation coefficient indicating the correlation between the target genes of a given ligand and the list of differentially expressed target genes in the “receiver” cell (**Figure 4B**). The receptors for the top predicted ligands *ANXA1, SPP1, TNF, CSF, CXCL9, IFNG, CD40LG, ITGB2, CD28, ICAM1, ADM*, and *IL18* all showed non-zero expression in pDCs. To identify which genes may be regulated by top-ranked ligands, putative ligand-gene interactions were scored by NicheNet according to “regulatory potential”, a graph-based likelihood for a ligand to regulate a particular target gene [37]. We next sought to identify candidate “sender cells” expressing these ligands (**Figure S4A**). We found that MPs and neutrophils expressed transcripts for pDC ligands, and we chose to further investigate these relationships. Ligand-mediated cell-cell interactions between MPs, neutrophils, and pDCs are visualized in the network diagram in **Figure 4B**. As shown, the differential expression of *AREG* is potentially regulated by IL-6 and SPP1 from MPs, and IL1-β from neutrophils, although the receptors for IL-6 and IL1-β could not be verified at the transcript level in pDCs in our dataset.

Additionally, given the enrichment of oncogenic pathways in epithelial cells of the BALF, we identified cell types and signaling molecules potentially regulating epithelial DEGs. We applied the NicheNet procedure outlined above for pDCs to club and ciliated cells of the BALF, resulting in the ligand-gene regulatory matrix shown in **Figure 4C**. The ligands *IL1B, IL1A, TNF, IFNG, APOE, IL1RN, OSM, EDN1, LIF, MIF, CSF2, SPP1, CD28, CCL2, VEGFA,* and *CCL5* show potential for regulating differential expression, and receptors for all of these ligands were manually verified at the transcript level in club or ciliated cells. Interestingly, *FOS* and *JUN* are among the genes potentially driven by the proposed ligands. As MPs and neutrophils appear to express the ligands of interest (**Figure S4B**), we investigated the role of these cells in regulating epithelial gene expression, visualized in the network diagram in **Figure 4C** (at larger scale in **Figure S4E**). As shown, MPs may induce both *FOS* and *JUN* through IL1-α, while neutrophils may induce both *FOS* and *JUN* through IL1-β, TNF, and IFNG, with additional regulation of *JUN* by EDN1.

Finally, we aggregated ligand-receptor relationships across all of the cell types in the BALF and proposed broadly-acting “pan-ligands” possibly contributing to the induction of transcriptional programs in the lung microenvironment. Following the procedure described above **(Figure 4A)** using each cell type in the BALF as a receiver, we identified a list of potential pan-ligands and developed a ligand-receiver cell correlation matrix (**Figure S4C**). This matrix was subsequently filtered to preserve ligands acting on over one-third of cell types in the BALF (≥6 cell types) with >10% receptor expression and thresholded Pearson correlation values. Of these candidates, we selected ligands differentially expressed between severe and moderate disease in one or more BALF cell types (**Figure S4D**) to further filter the ligand-receiver matrix. Interestingly, the *IL6* transcript was not found to be differentially expressed in any of the BALF cell types and failed at this filtering step. After filtering, we arrived at a list of differentially expressed pan-ligands implicated in broad severity-specific responses across cell types in the lung, consisting of *IFNG, IL18, CCL3, CCL8*, and *SPP1*. These ligands were differentially expressed in at least one BALF sender cell type and act on over one-third of the cell types in the BALF (**Figure 4D**), indicating these ligands should be further investigated to elucidate their role in the life-threatening immunopathology of severe COVID-19.

## DISCUSSION

In the course of a severe SARS-CoV-2 viral infection, acute pulmonary damage has been observed concomitantly with elevated cytokine levels in serum and infiltration of macrophages and lymphocytes into multiple organs, indicative of both local and systemic immunological responses [11, 15]. To better understand these dynamics, we leveraged scRNA-seq data from BALF and blood to compare severity-specific shifts in cellular composition at the local and systemic levels. In BALF, compositional analysis indicated expansion of NK cells, T cells (general T cell population, CD4+ naive T cells, and CD4+ effector Treg cells), and pDCs in moderate patients compared to both severe and control patients. Although severity-specific compositional changes in BALF samples remain largely unaddressed in the literature [27], the severity-specific lymphopenia of NK cells and CD4+ naive T cells that we report in BALF agrees with findings from compositional studies of PBMCs [30, 31, 34]. We additionally report severity-specific compositional shifts in lung MP populations and identify clusters expressing *MRC1*, *C1QA*, and *FABP4* suggesting an M2 macrophage-like phenotype expanded in control subjects. Moderate donors showed expanded clusters expressing *HLADQA1* and *HLADQA2*, indicating increased antigen presentation through HLA class II. Expanded clusters in severe donors expressing *SPP1* as well as various chemokines including *CCL2*, *CCL3*, and *CCL8* are likely associated with an M1 macrophage-like phenotype, in agreement with previous analyses [27]. Strong macrophage expression of osteopontin, the gene product of *SPP1,* has been previously observed during influenza A infection, suggesting that macrophages in severe COVID-19 may be following a similar viral response to influenza [43]. Compositional changes were not readily apparent in PBMCs when comparing ARDS patients to non-ARDS patients, although we observed an expansion of multiple plasma cell types in ARDS patients compared to healthy control subjects, consistent with previous reports [23, 33, 34, 44].

Despite recent advances, the immunological signatures of severe COVID-19 remain largely uncharacterized in both the lung [27, 28] and systemic circulation [34, 36]. In particular, a need exists to define markers of severity that can be used to assess patient disease trajectory in a clinical setting for the purposes of triage or to inform the use of immunomodulatory treatment strategies [34]. Here, we leveraged BALF and PBMC transcriptional data to nominate cell surface proteins that could be incorporated into a flow cytometry panel for COVID-19 patient immunophenotyping. Notably, in BALF samples we observed upregulation of *AREG* in pDCs and mDCs of patients undergoing a severe disease course compared to patients who experienced moderate disease. The *AREG* transcript was verified to be expressed by dendritic cells in bulk RNA-seq data [39, 40], has been previously shown to orchestrate tissue homeostasis during influenza infection [45, 46], and has been previously labeled in flow cytometry studies [47]. Additionally, *TIMP1*, *VAMP5,* and *IL1R2* were strongly differentially expressed in BALF MPs during severe disease and have been previously documented to play a role in the immune response to respiratory viral infections. *TIMP1* has been shown to promote deleterious immune responses in the lung during murine influenza infection [48], *VAMP5* is implicated in signaling pathways regulating influenza viral replication *in vitro* [48, 49], and *IL1R2* encodes a decoy receptor for IL1-β that is upregulated during severe influenza [50]. All of the BALF identified surface markers in **Figure 2C** with the exception of *VAMP5*, *HLA-DRA*, and *CCR7* were specifically differentially expressed between severe and moderate donors but not between moderate and control donors (**Figure S2B**). Investigating cell surface markers in PBMCs, we found that *IL7R* on NK cells as well as *S1PR1* (*CD363*) on γδ T cells differentiate ARDS from non-ARDS COVID-19 blood samples. IL-7 stimulation through IL-7R has been shown to promote NK cell survival by inhibiting apoptosis [50, 51]. *S1PR1* is a particularly interesting marker due to previous literature documenting its role in stimulating highly inflammatory pathways such as JAK/STAT and mTOR/PI3K/Akt. Additionally, suppression of the S1PR1 cognate ligand S1P1 decreased mortality rates during influenza infection in mice [41, 42, 52]. All of the identified PBMC surface markers in **Figure 2F** were specifically differentially expressed between ARDS and non-ARDS donors but not between non-ARDS and control donors (**Figure S2E**). Our investigation revealed a small number of common surface markers of interest between BALF and PBMC samples, namely *HLADQA1* and *HLADQA2* on MPs/CD16+ monocytes and *HLADRA* on MPs/CD14+ monocytes; these HLA class II transcripts are downregulated with increasing severity level in both compartments. Similar observations of a severity-specific loss of HLA-DR on CD14+ monocytes and T cells have been previously observed in recent scRNA-seq and flow cytometry studies [29, 30, 31, 32, 34, 36]. Together, these findings suggest transcripts that should be studied further at the protein level as potential markers of a severe or ARDS disease course for immunophenotyping patients through BALF sampling or a convenient blood draw.

Cell type-specific immunopathological responses in severe COVID-19 have been investigated at both local and systemic levels in recent studies comparing severe patients to healthy controls [21, 22, 23, 24, 25, 26, 32, 53, 54, 55]. We sought to build on this knowledge by determining the cell-type specific biological responses that differentiate more finely stratified disease severity levels in the lung and systemic circulation. In myeloid and lymphoid cells of the BALF, we observed a concerted enrichment of the TNF-α signaling via NF-κB pathway in severe compared to moderate donors, indicating the upregulated expression of factors such as *NFKBIA* and *TNFAIP3* in severe disease. We further noted IL-2 and IL-6 signaling pathway enrichment in cell types including MPs, mast cells, and neutrophils. *IL2RA* expression was observed in mast cells, *IL6R* was expressed in MPs, mast cells and neutrophils, and neither the *IL2* nor *IL6* transcripts were well-captured in the dataset. IFN-α (type I) and IFN-γ (type II) pathway responses showed mixed enrichment and depletion, with type I and II interferon signaling apparently decreased in a majority of BALF T cell subtypes. These data are consistent with previous observations of low expression of *IFNG* and *TNF* in cytotoxic T lymphocytes of BALF derived from severe patients [28]. MPs in the BALF demonstrated a nearly homogenous enrichment of the IL-6, IL-2, and TNF-α signaling via NF-κB pathways across the diversity of clusters. When we compared disease strata based on ARDS diagnosis in PBMCs, fewer pathways demonstrate enrichment with increasing disease severity. Nevertheless, a striking depletion of the type II interferon response was observed in nearly every cell type studied, along with a broad depletion of the type I interferon response and TNF-α signaling via NF-κB pathways (including downregulation of *TNFAIP3* and *NFKBIA*). This result corroborates a previously reported correlation between suppressed type I and II interferon responses and disease severity in a study of PBMCs that was not specific to cell type [33], as well as extending findings of increased interferon response in monocytes during moderate disease but not severe disease [34, 36]. Our findings do not preclude the observation of increased TNF-α signaling in the bloodstream of severe and critical COVID-19 patients [33], as *TNFAIP3* depletion has been shown to cause increased TNF-α expression [56].

We next examined the similarity of pathway-level changes across the severity stratifications provided in the BALF and PBMC datasets. Comparing BALF and PBMC responses to increasing disease severity level in consanguineous cell types showed shared suppression of the type I interferon response in CD4+ naive, CD8+ effector, and γδ T cells and of the type II interferon response in mature B, proliferating plasma, CD4+ naive, CD8+ effector, and γδ T cells. For other cell types including MPs/monocytes, NK and CD8+ effector T cells, the type II interferon response appears to be increased in cells of the lung but decreased in the blood. Moreover, our analysis identifies divergent severity-specific responses in the BALF and blood in terms of the TNF-α signaling via NF-κB pathway for nearly all cell types. Notably, the concerted TNF-α/NF-κB pathway enrichment across BALF cell types is indicative of a strong upregulation of *TNFAIP3* and *NFKBIA.* Conversely, the depleted pathway across PBMC cell types corresponds to *TNFAIP3* and *NFKBIA* downregulation. The A20 protein encoded by *TNFAIP3* is an NF-κB inhibitor that functions as a “brake” on antiviral signaling and inflammatory responses [56, 57, 58, 59] whose deletion in mouse models improved survival during influenza infection [59]. *TFNAIP3* differential expression was largely co-directional with *NFKBIA* (coding for IκBα), a gene whose upregulation is a key feature of the human blood response to SARS-CoV as well as the lung response to SARS-CoV and MERS-CoV infection [60]. Our results suggest that the A20/IκBα axis likely plays a role in SARS-CoV-2 infection as well, with *TNFAIP3* inhibiting NF-κB-mediated antiviral responses during severe disease at the site of local infection yet promoting a systemic inflammatory response through its absence during ARDS in the periphery. We note in this compartmental comparison that the BALF and PBMC clinical stratifications represent different parts of the disease severity spectrum. Therefore, we interpret differences in compartmental responses with respect to relative levels of clinical severity rather than absolute severity stages. Taken together, our data suggest that transcriptional programs are differentially induced with increasing COVID-19 severity, while the specific responses are nuanced according to cell type and local versus systemic immune environment.

In epithelial cells of the BALF, we detected the surprising upregulation of *FOS* and *JUN*, along with the downregulation of *C2orf40* in severe compared to moderate donors. Due to the involvement of these genes in oncogenic programs, we further investigated pathway-level alterations in epithelial cells and detected the enrichment of epithelial-to-mesenchymal transition, K-Ras, PI3K/AKT and p53 pathways. We note that a separate analysis of epithelial cells from the same BALF dataset failed to detect differential enrichment of these pathways through GSEA when comparing control samples to pooled moderate and severe samples [21, 53], likely as a result of this pooling and using a much shorter list of statistically tested genes (a single set of 118 genes common to all epithelial cells compared to sets of 1,561 genes for ciliated cells and 1,092 genes for club cells). Our results may be cautiously interpreted as providing evidence for the molecular underpinnings of the morphological changes known to occur in the severely damaged lung epithelia of severe COVID-19 patients, previously described as cellular proliferation resembling atypical adenomatous hyperplasia, *in situ* adenocarcinoma, or even invasive adenocarcinoma [15].

Following the observation of pathways enriched in the epithelia of severe COVID-19 patients, we employed ligand-receptor network analysis to nominate potential immune-epithelial communication networks that may explain transcriptional differences between severe and moderate disease courses. We identified 16 ligands implicated in driving the differential expression in ciliated and club cells, of which *IL1B, TNF*, and *IFNG* are predicted to regulate *FOS* and *JUN*, with *IL1A* additionally regulating *JUN.* Exploring the expression of these 16 ligands across the diversity of BALF cell types indicates that MPs and neutrophils may act as “sender cells” signaling to the epithelium, as suggested in recent reports [21, 28, 53]. In particular, aberrant neutrophil responses have been implicated in severe COVID-19 through a number of studies focusing on either the lung or the blood [21, 28, 34, 35, 36, 53]. On further analysis, we observed potential MP signaling through *CCL2, SPP1, CCL13, APOE, INHBA,* and *IL1A*, and neutrophil signaling through *CCL3, CCL5, EDN1, IFNG, TNF, IL1RN*, and *IL1B*, among others. These ligands may represent key players in immune-to-epithelial signaling networks distinguishing moderate from severe disease progression in the lung microenvironment during SARS-CoV-2 infection.

Our analysis of the transcriptional regulation of cell surface proteins implicated *AREG* as a potential marker unique to severe disease in pDCs. This finding appears consistent with previous literature showing that amphiregulin aids in the maintenance of epithelial integrity and tissue repair during infection or injury, [45, 61] as patients with severe COVID-19 experience extreme pulmonary damage. Expanding on this knowledge, we utilized ligand-receptor network analysis to provide orthogonal information suggesting how pDC differentially expressed genes, including *AREG*, may be regulated through soluble or cell-surface mediators. NicheNet revealed that MPs and neutrophils appear to signal to pDCs through *SPP1*/*IL6* and *IL1B* respectively to regulate the expression of *AREG* as well as various other genes. *SPP1* in particular appears to be better-evidenced than *IL6* and *IL1B* as its active receptors *CD44* and *ITGB1* are present on over 20% of pDCs in severe donors. Our network analysis suggests that the expansion of a population of *SPP1*-expressing MPs during severe disease, observed here and in Liao et al. [27], may partly account for the upregulation of AREG by pDCs. We recommend further study of the potential for *AREG* induction by SPP1 in addition to investigation of AREG surface expression as a marker of severe disease.

Immune blockades, including the IL-6R antagonist Tocilizumab, are a class of treatments currently under clinical use and research for COVID-19 intended to dampen the hyperactive immune response by targeting key nodes in a signaling network [23, 30, 31]. Here, we sought to utilize ligand-receptor network analysis to nominate ligands as potential key regulators of severity-specific dysregulation in the lung microenvironment during severe COVID-19. Our analysis suggests *IFNG, IL18, CCL3, CCL8*, and *SPP1* as candidate “pan-ligands” that may induce transcriptional regulation in over one-third of cell types identified in the BALF during severe disease. Notably, we did not identify *IL6* as a potential pan-ligand as its differential expression between moderate and severe patients in our study was not statistically significant in any cell type. All of the pan-ligands we report have been previously implicated in SARSr-CoV infection, with IL-18, CCL8, and IFN-γ having been detected in serum from COVID-19 patients [9, 13, 14], *SPP1* upregulation measured in microarrays from SARS-CoV-infected nonhuman primates, and *CCL3, CCL8*, *IL18,* and *IFNG* upregulation detected in single-cell or bulk RNA-seq from the lungs of COVID-19 patients [21, 22, 27, 28]. Importantly, our data supports accumulating evidence for a nuanced interferon response in COVID-19 depending on disease severity, cell type, and local versus systemic immune environment, as discussed above [25, 26, 28, 33, 34, 36, 55]. We suspect that treatment strategies consisting of a direct type II interferon blockade (NIH clinical trial NCT04324021) [62, 63] could exert different effects systemically and locally depending on the patient’s clinical condition when administered. Taken together with previous findings, our data suggests *IFNG, IL18, CCL3, CCL8*, and *SPP1* as candidate targets for the treatment of severe COVID-19 that warrant further study.

In summary, our findings emphasize unique roles for the cells of the lung microenvironment and systemic circulation in the immunopathology of ARDS and severe COVID-19. We recommend further investigation of differentially expressed cell surface markers to determine their utility in immunophenotyping patients according to suspected disease course to aid in triage or to inform optimal treatment options. Our transcriptional analyses show that pathway enrichment differs between cells of the lung and blood, with concerted immunological responses within each compartment mirroring viral respiratory diseases including influenza, SARS, and MERS. Finally, we nominate a small number of broadly-acting ligands as potential drivers of severity-specific transcriptional regulation in the severely damaged lung microenvironment of COVID-19 patients.

## METHODS

### Data Acquisition

All single cell RNA-sequencing (scRNA-seq) data used in this analysis were obtained from publicly available datasets. BALF scRNA-seq gene-barcode matrices from 9 COVID-19 patients (6 severe and 3 moderate), and 3 healthy control subjects were obtained from the GEO under accession number GSE145926. [27] ScRNA-seq data from an additional healthy BALF donor was obtained from the GEO under accession number GSM3660650. [27, 64] Severity-level stratifications for BALF donors (severe/moderate/healthy) were used exactly as provided in the original manuscript. 5 out of 6 severe BALF donors were invasively ventilated, with the exception of S1. [27] PBMC scRNA-seq gene-barcode matrices from 7 patients (4 samples from patients with ARDS and 4 samples from patients without ARDS, where one patient who was sampled twice fell contributed to both groups at different stages of the disease course) and 6 healthy control donors were obtained from the GEO under accession number GSE150728. Severity-level stratifications of PBMC donors (ARDS/non-ARDS/healthy) were implemented using the exact ARDS classification provided in the original manuscript based on the Berlin criteria. One donor in the PBMC dataset was sampled in the non-ARDS stage (M4) and re-sampled after the onset of ARDS (S7). All ARDS patients in the PBMC dataset required invasive ventilation. [32]

### Data Preprocessing and Multi-Donor Integration

Gene-barcode matrices obtained from the GEO were preprocessed using Seurat (v. 3.1.5) in R (v. 4.0.0). Matrices were filtered to preserve cells with a UMI count over 1,000, gene count between 200 and 6,000, and expression of less than 10% mitochondrial RNA. For PBMC data, cells were additionally filtered to preserve cells expressing less than 20% 18S RNA and less than 20% 28S RNA. Gene-barcode matrices unique to each donor were subsequently normalized using the ‘NormalizeData’ function and the 2,000 most highly variable genes (features) were identified via the ‘FindVariableFeatures’ function using a variance stabilizing transformation. Individual donor datasets were subsequently aligned and integrated using the standard Seurat v3 multi-donor integration workflow, finding pairwise anchors using the ‘FindIntegrationAnchors’ function acting on 50 dimensions with default parameters (k.filter=200), then applying the ‘IntegrateData’ function on 50 dimensions. Multi-donor integration and all further analyses were performed separately on BALF and PBMC datasets. Following integration, “raw” RNA expression values were normalized using log-normalization via the ‘NormalizeData’ function and the 2,000 most variable features were identified using ‘FindVariableFeatures’. Raw expression levels were left unscaled so that scaling could later be applied as needed (e.g. for generating marker gene heatmaps). In parallel, “integrated” expression values were scaled using the ‘ScaleData’ function, regressing out UMI count, number of genes, and percent rRNA. PBMC data contained rRNA transcripts to regress, whereas BALF data did not.

### Dimensionality Reduction and Clustering

Dimensionality reduction via principal component analysis (PCA) was carried out using the ‘RunPCA’ function and the first 100 principal components were retained. This reduction was projected onto the entire dataset prior to clustering. Clustering was performed by finding nearest neighbors using the ‘FindNeighbors’ function in Seurat acting on 50 dimensions, then running the Louvain algorithm via the ‘FindClusters’ function with a resolution parameter of 1.4. The resulting clustered datasets for BALF and PBMC were separately visualized by uniform manifold approximation and projection (UMAP) acting on the first 50 principal components.

### Cell Type Annotation and Iterative Data Integration

Cell type annotation was conducted using the resultant clusters for BALF and PBMC data. Clusters were annotated according to the average expression levels of canonical marker genes identified in the original papers for the BALF and PBMC datasets [27, 32], the broader literature, and the Human Protein Atlas [65]. Clusters deemed identical by similar presence of marker genes were merged during annotation. A list of the marker genes used for cell type annotation is available in Supplemental Figure S1, along with a visualization of expression levels. Following cell type annotation for the BALF, populations annotated as mononuclear phagocytes (consisting of one merged cluster) and T/NK cells (consisting of clusters annotated as “NK”, “Proliferating T” and “Mixed T” cells) were separately re-integrated to enable clustering at a finer resolution. For mononuclear phagocytes, iterative data integration was performed using the integration procedure described above, with a k.filter parameter of 112 for ‘FindIntegrationAnchors’ and a resolution parameter of 0.8 for ‘FindClusters’. The resulting fine-scale mononuclear phagocyte clusters were left unannotated during further analysis, but were tested using the BALF marker gene panel to identify probable doublets. For T and NK cells, iterative data integration was performed using the integration procedure described above, with a k.filter parameter of 175 and a resolution parameter of 0.8. In order to filter anchors for T/NK re-integration, one healthy control donor (HC2, 60 cells) and one severe donor (S3, 72 cells) with small numbers of T/NK cells were excluded from the integration and subsequent analysis. T and NK cell fine-level clusters were annotated according to a panel of canonical T and NK cell marker genes, available in Supplemental Figure S1. The original integration of PBMC mutli-donor datasets allowed cell type annotation at a fine level and did not require iterative data integration.

### Compositional Analysis

Compositional changes across disease severity categories were assessed for BALF and PBMC datasets separately using: 1) donor-specific fractional contributions to each cell type in the total pool of cells recovered from all donors, 2) donor-specific percentages of cell types comprising each pool of cells recovered from individual donor severity conditions (severe, moderate, or healthy control for BALF, ARDS, non-ARDS, or healthy control for PBMCs), and 3) percentages of cell types in each donor sample z-scored across donors. Here, we did not employ low-powered statistical tests but instead sought to use method 3 to highlight donor-to-donor variation in cell type composition and identify robust trends across donors indicating possible expansion or cytopenia.

### Differential Expression Analysis

Differential expression analysis was performed in Seurat using the ‘FindMarkers’ function utilizing model-based analysis of single-cell transcriptomics (MAST) statistical framework through the “MAST” R package (v. 3.11) [66]. Differential expression analysis was conducted for all cell types in the integrated BALF dataset across three permutations of donor groups: severe vs. moderate, moderate vs. control, and severe vs. control. For all cell types in the PBMC dataset, differential expression analysis was conducted across three permutations of donor groups: ARDS vs. non-ARDS, non-ARDS vs. control, and ARDS vs. control Significantly differentially expressed genes (DEGs) are indicated by having a MAST adjusted p<0.05 and a natural log fold change threshold of 0.25 was applied.

### Surface Marker Identification

The identification of cell surface markers indicative of severe disease was performed by cross-referencing all DEGs across disease severity levels in the integrated BALF and PBMC datasets with entries in the Cell Surface Protein Atlas (CSPA) [38]. DEGs were considered to be differentially expressed surface markers if they were included in the “high confidence” CSPA category and showed significant differential expression (adjusted p<0.05) with an absolute value average natural log fold change greater than 0.693 (or log2 fold change >1). Differentially expressed surface markers were further analyzed to verify low expression in moderate or non-ARDS and control donor samples. Finally, differentially expressed surface markers were verified to be expressed in the cell type indicated by our data by cross referencing with the Immune Cell Atlas human bulk RNA-seq data [39, 40].

### Gene Set Enrichment Analysis (GSEA)

Gene set enrichment analysis (GSEA) [67] was performed using the “fgsea” package (v. 1.12.0) [68] in R. Differentially expressed gene lists generated using MAST were ranked by −log_10_(p) multiplied by the sign of the average natural log change, as previously demonstrated in Debski et al. and Riemand et al. [69, 70], using the MAST adjusted p-value. Average natural log fold change was used to break ranking ties and this value also served as an input to GSEA to quantify the correlation between the gene and phenotype, as −log_10_(p) values become arbitrarily large. GSEA results were interpreted according to normalized enrichment score (NES) and an adjusted p-value, with a p<0.05 significance threshold. Pathways with positive NES were defined throughout the text as “enriched” and pathways with negative NES were defined as “depleted”.

### Gene Ontology (GO) and Transcription Factor (TF) Analysis

Further analysis of significantly differentially expressed genes was performed using the “EnrichR” (v. 2.1) package in R. Gene ontology (GO) analysis was conducted on the DEGs between the severe and moderate samples for both the BALF and PBMC, evaluating the GO “biological function” annotations representing large scale biological programs. Transcription factor (TF) analysis was similarly conducted on the DEG list to query the regulation of TFs identified by the ENCODE and ChEA ChIP-X databases [71, 72]. GO and TF results were ranked based on the EnrichR ‘Combined Score’ metric and significance was determined using an adjusted p-value threshold of p<0.05.

### Ligand-Receptor Network Analysis for BALF Cells

To investigate the intracellular interactions potentially contributing to the observed differential gene expression between the severe and moderate sample populations in the BALF, we employed the ligand-receptor interaction tool NicheNet via the “nichenetr” package (v. 0.1.0) in R. [37] Differentially expressed target genes between severe and moderate disease in a “receiver” cell population of interest were identified using the ‘FindMarkers’ function in Seurat with criteria of p<0.05, average natural log fold change >0.25, and expression in over 10% of the receiver cells. Concurrently, a list of potential receptors expressed in over 10% of cells in the severe disease receiver population was generated using NichNet. A list of “sender” cells was created comprising all cell types in the BALF, including the receiver cell type to account for the possibility of autocrine signaling. For each sender cell population, potential ligands were inferred using the NichNet ligand-receptor network applied to genes expressed in over 10% of the severe disease sender population. Ligand activities were predicted using the ‘predict_ligand_activities’ function in NicheNet, and the top 20 ligands were selected by Pearson correlation coefficient. Differentially expressed target genes ranking among the 100 most strongly predicted targets of the top 20 ligands were given a “regulatory potential” interaction score. The upper 50% of targets by regulatory potential were visualized for plasmacytoid dendritic cells (pDCs), and the upper 15% of targets were visualized for epithelial cells (ciliated cells and club cells). To specifically analyze signaling from mononuclear phagocytes and neutrophils, differentially expressed target genes ranking among the 250 most strongly predicted targets of the top 20 ligands were used, with the upper 25% of these targets according to regulatory potential visualized in circos plots. Non-signaling molecules and molecules acting on non-coding RNA targets were manually removed. Receptors for the top 20 ligands were identified using the CellPhoneDB web server [73], and non-zero expression was verified in the severe disease receiver cell population of interest using Seurat.

Identification of potential broadly acting “pan-ligands” was conducted by replicating the above NicheNet procedure iteratively using each cell type of the BALF as a receiver. For each receiver cell type, ligands with positive Pearson correlation were filtered based on 1) either belonging to the list of top 20 ligands or having a Pearson correlation greater than 0.1, and 2) presence of the corresponding receptor in over 10% of cells (manually validated using CellPhoneDB [73]). Only ligands meeting these criteria for over one-third of cell types in the BALF were preserved for further analysis. Of these, the ligands differentially expressed in severe disease compared to moderate disease in at least one BALF cell type were classified as potential pan-ligands.

### Statistical Analysis

Differential gene expression was analyzed using the “MAST” package statistical framework through the ‘FindMarkers’ function in Seurat. Differential expression statistical significance was qualified using the MAST false discovery rate (FDR) adjusted p-value, with a significance threshold of p<0.05. Pathway enrichment was analyzed using the standard GSEA method in the “fgsea” package. Statistical significance of normalized enrichment scores was qualified using the fgsea FDR adjusted p-value, with a significance threshold of p<0.05. Differential gene expression relevant to ligand-receptor interactions was evaluated using the NichNet pipeline implementing the Wilcoxon rank sum test through the ‘FindMarkers’ function in Seurat. Statistical significance was qualified using the ‘FindMarkers’ Bonferroni-adjusted p-value, with a significance threshold of p<0.05. GO and TF analysis was conducted using the “EnrichR” package, and statistical significance was qualified using the EnrichR adjusted p-value based on Fisher’s exact test, with a significance threshold of p<0.05.

## Data and Code Availability

All data used in the study are available in the GEO under accession numbers GSE145926, GSM3660650, and GSE150728. All code used for analysis will be made available in a public repository.

## ACKNOWLEDGEMENTS

The authors gratefully acknowledge Dr. Paul Blainey, Cal Gunnarsson, Bianca Lepe, Megan Tse, Andy Kim, and Harvey Yang for providing early-phase feedback. Figures were partially created using BioRender.com. The authors thank Jon Henninger for input on figure design. K.O. is funded by the National Science Foundation Graduate Research Fellowship Program. The authors would like to thank the MIT Department of Biological Engineering for additional support throughout the research effort.

## AUTHOR CONTRIBUTIONS

K.O. and J.K. conceived of the ideas as a course project for 20.440 at MIT then undertook an independent research effort. K.O. and J.K. contributed equally to the analysis and sought guidance and supervision from B.B. throughout the research process. K.O., J.K., and B.B. wrote and edited the manuscript.

