## Supplemental Information for "Dissecting the common and compartment-specific features of COVID-19 severity in the lung and periphery with single-cell resolution"

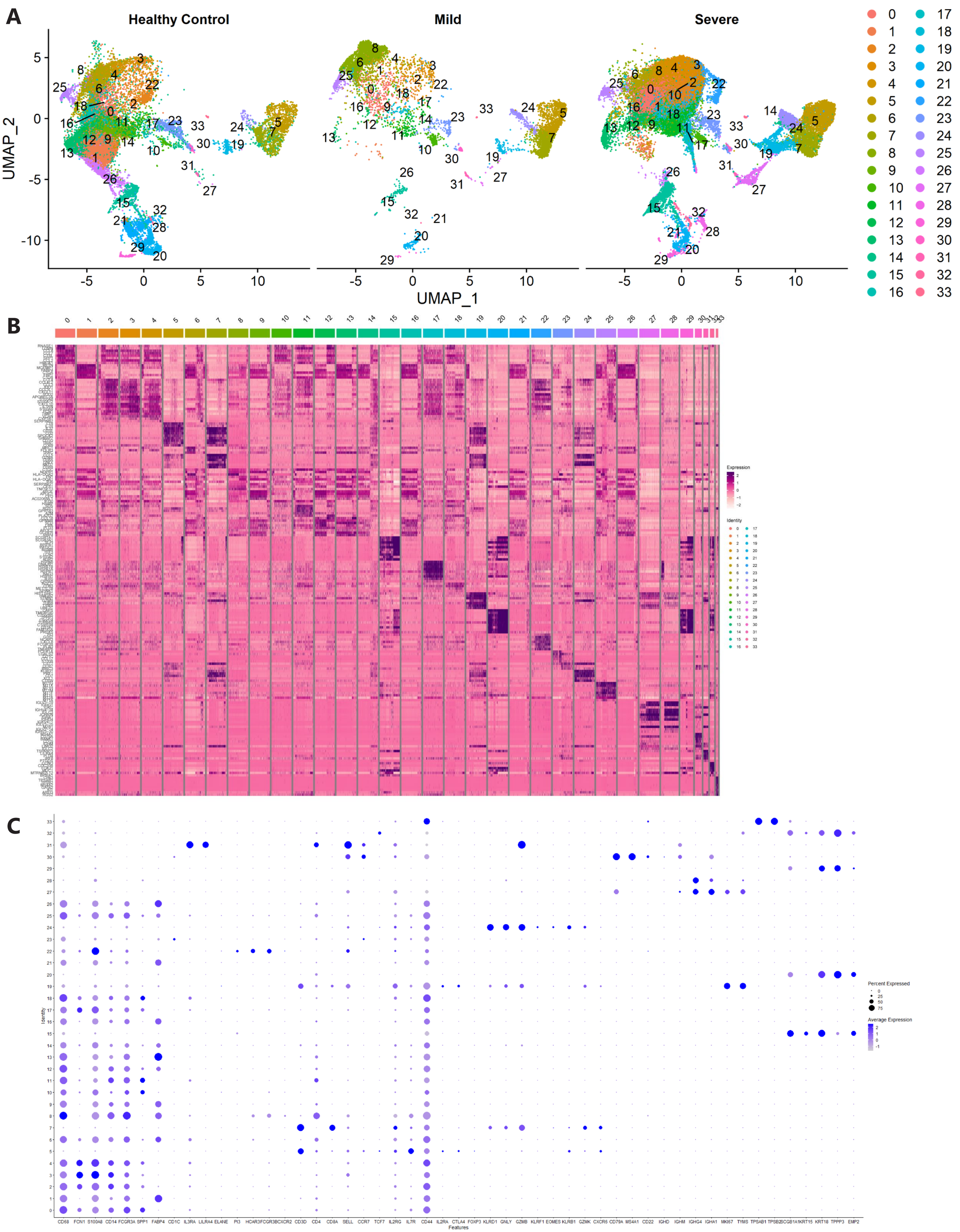

A
Healthy Control
Mild
Severe
UMAP_1
UMAP_2
B
Expression
Identity
C
Percent Expressed
Average Expression
Identity
Features
CD68 FCN1 S100A8 CD14 FCGR3A SPP1 FABP4 CD1C IL3RA LILRA4 ELANE PI3 HCAR3 FCGR3B CXCR2 CD3D CD4 CD8A SELL CCR7 TCF7 IL2RG IL7R CD44 IL2RA CTLA4 FOXP3 KLRD1 GNLY GZMB KLRF1 EOMES KLRB1 GZMK CXCR6 CD79A MS4A1 CD22 IGHD IGHM IGHG4 IGHA1 MKI67 TYMS TPSAB1 TPSB2 SCGB1A1 KRT15 KRT18 TPPP3 EMP2

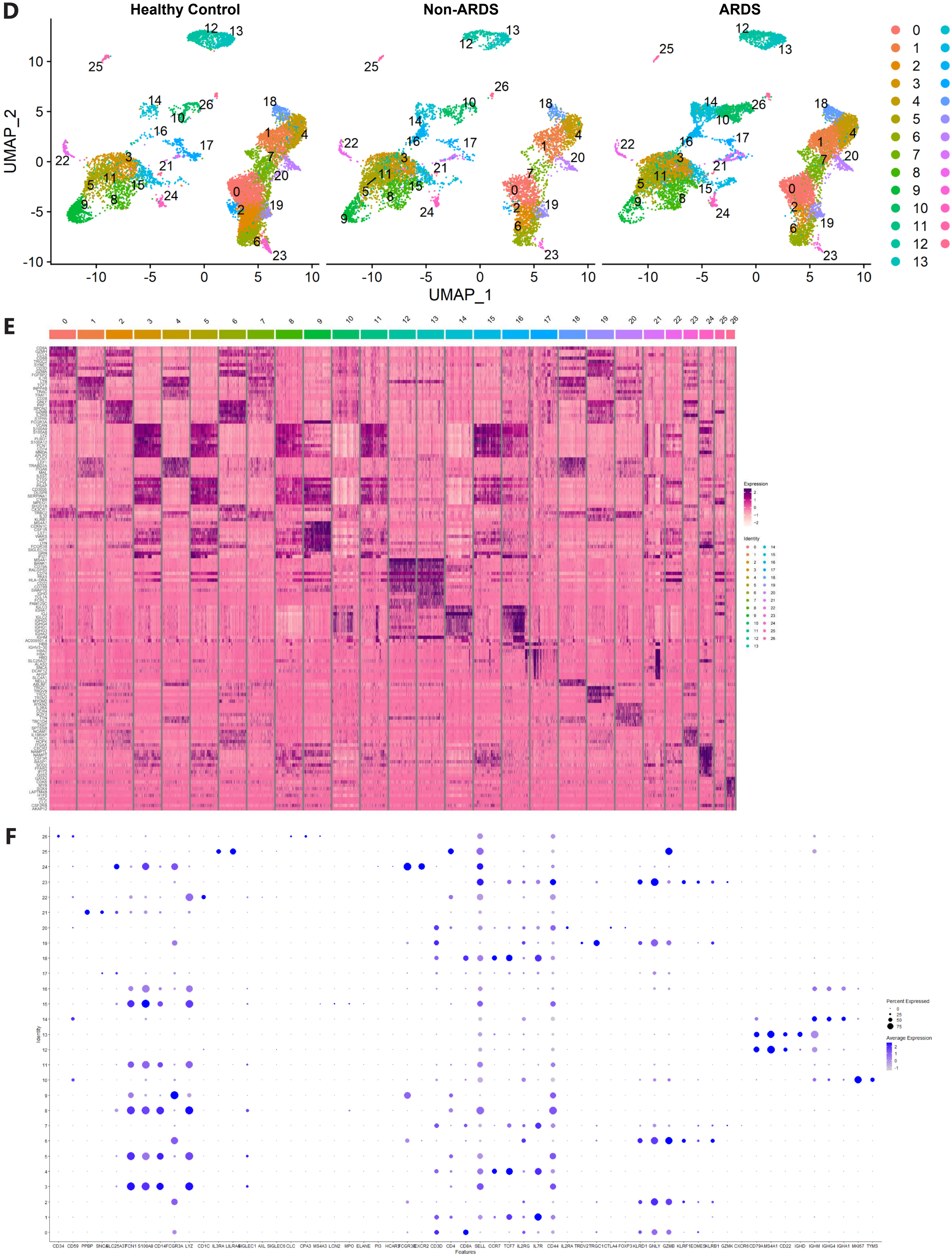
D
Healthy Control
Non-ARDS
ARDS
UMAP_2
UMAP_1
E
Expression
Identity
F
Percent Expressed
Average Expression
Identity
Features
CD34 CD59 PPBP SNCA SLC25A37 FCN1 S100A8 CD14 FCGR3A LYZ CD1C IL3RA LILRA4 SIGLEC1 AXL SIGLEC6 CLC CPA3 MS4A3 LCN2 MPO ELANE PI3 HCAR3 FCGR3B CXCR2 CD3D CD4 CD8A SELL CCR7 TCF7 IL2RG IL7R CD44 IL2RA TRDV2 TRGC1 CTLA4 FOXP3 KLRD1 GNLY GZMB KLRF1 EOMES KLRB1 GZMK CXCR6 CD79A MS4A1 CD22 IGHD IGHM IGHG4 IGHA1 MKI67 TYMS

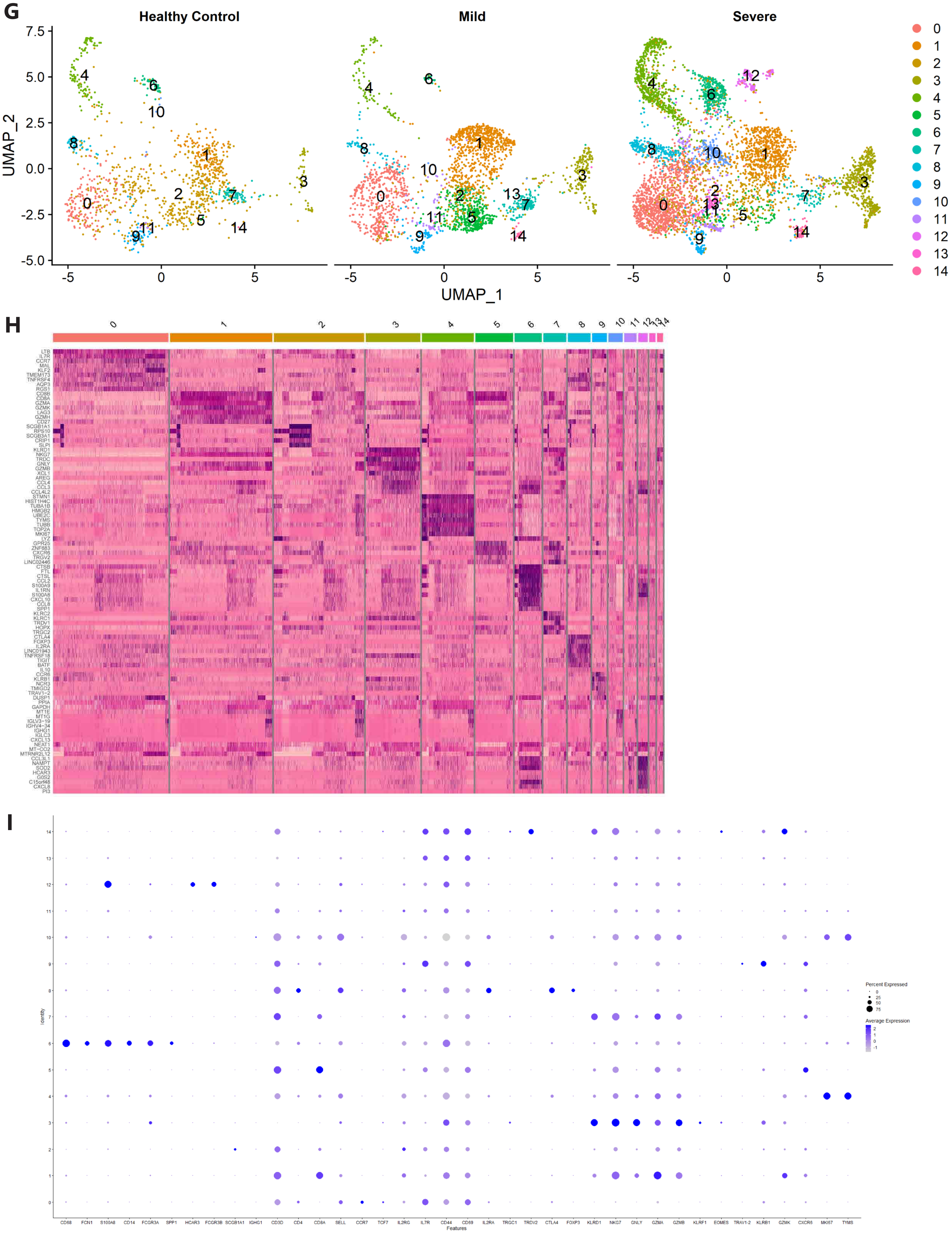
G
Healthy Control
Mild
Severe
UMAP_2
UMAP_1
7.5
5.0
2.5
0.0
-2.5
-5.0
-5
0
5
H
LTB
IL7R
CCR7
MAL
KLF2
TMEM173
TNFRSF4
AQP3
RGS1
CD8B
CD8A
GZMA
GZMK
LAG3
GZMH
CD27
SCGB1A1
RPS10
SCGB3A1
CRIP1
SLPI
KLRD1
NKG7
TRDC
GNLY
GZMB
XCL1
AREG
CCL4
CCL3
CCL4L2
STMN1
HIST1H4C
TUBA1B
HMGB2
UBE2C
TYMS
TUBB
TOP2A
MKI67
LYZ
GPR25
ZNF683
CXCR6
TRGV2
LINC02446
CTSB
FTL
CTSL
CCL2
S100A9
IL1RN
S100A8
CXCL10
CCL8
SPP1
KLRC2
KLRC1
TRDV1
HOPX
TRGC2
CTLA4
FOXP3
IL2RA
LINC01943
TNFRSF18
TIGIT
BATF
IL10
CCR6
KLRB1
NCR3
TMIGD2
TRAV1-2
DUSP1
PPIA
GAPDH
MT1E
MT1G
IGLV3-19
IGHV4-34
IGHG1
IGLC3
CXCL13
NEAT1
MT-CO2
MTRNR2L12
CCL3L1
NAMPT
SOD2
HCAR3
G0S2
C15orf48
CXCL8
PI3
I
Identity
Percent Expressed
0
25
50
75
Average Expression
2
1
0
-1
CD68
FCN1
S100A8
CD14
FCGR3A
SPP1
HCAR3
FCGR3B
SCGB1A1
IGHG1
CD3D
CD4
CD8A
SELL
CCR7
TCF7
IL2RG
IL7R
CD44
CD69
IL2RA
TRGC1
TRDV2
CTLA4
FOXP3
KLRD1
NKG7
GNLY
GZMA
GZMB
KLRF1
EOMES
TRAV1-2
KLRB1
GZMK
CXCR6
MKI67
TYMS
Features

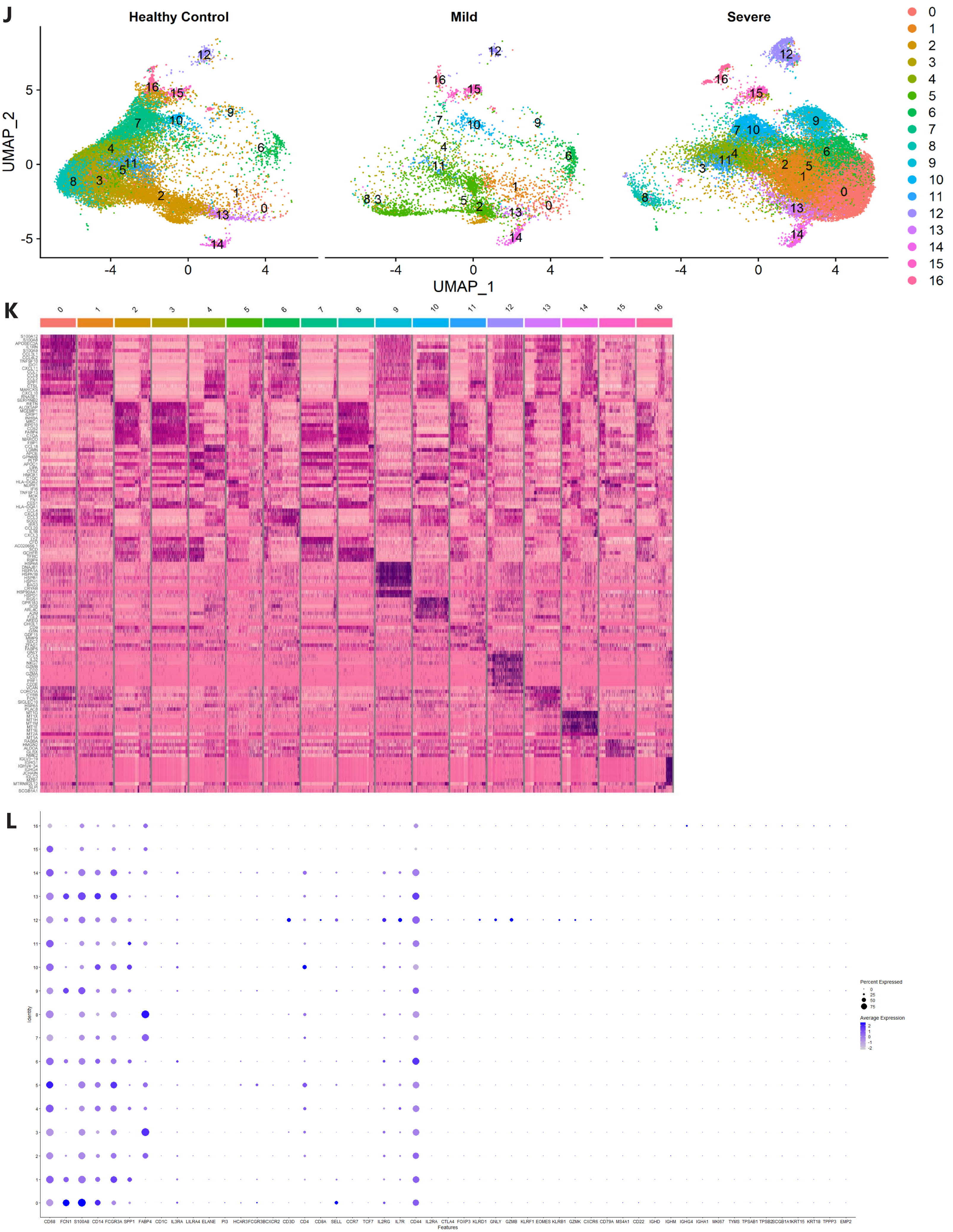
J
Healthy Control
Mild
Severe
UMAP_2
UMAP_1
K
L
Identity
Features
CD68 FCN1 S100A8 CD14 FCGR3A SPP1 FABP4 CD1C IL3RA LILRA4 ELANE PI3 HCAR3 FCGR3B CXCR2 CD3D CD4 CD8A SELL CCR7 TCF7 IL2RG IL7R CD44 IL2RA CTLA4 FOXP3 KLRD1 GNLY GZMB KLRF1 EOMES KLRB1 GZMK CXCR6 CD79A MS4A1 CD22 IGHD IGHM IGHG4 IGHA1 MKI67 TYMS TPSAB1 TPSB2 SCGB1A1 KRT15 KRT18 TPPP3 EMP2
Percent Expressed
Average Expression

M

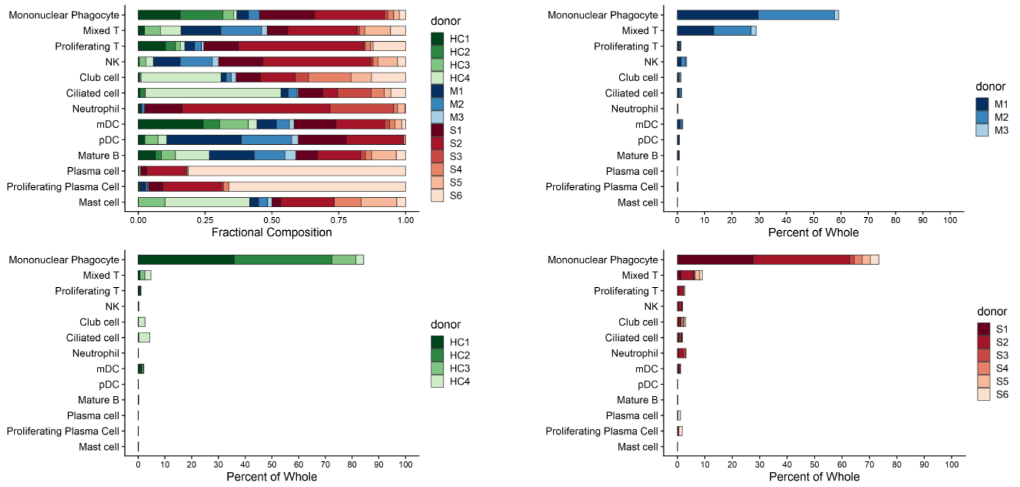
Mononuclear Phagocyte
Mixed T
Proliferating T
NK
Club cell
Ciliated cell
Neutrophil
mDC
pDC
Mature B
Plasma cell
Proliferating Plasma Cell
Mast cell
Fractional Composition
Percent of Whole
donor
HC1
HC2
HC3
HC4
M1
M2
M3
S1
S2
S3
S4
S5
S6

N

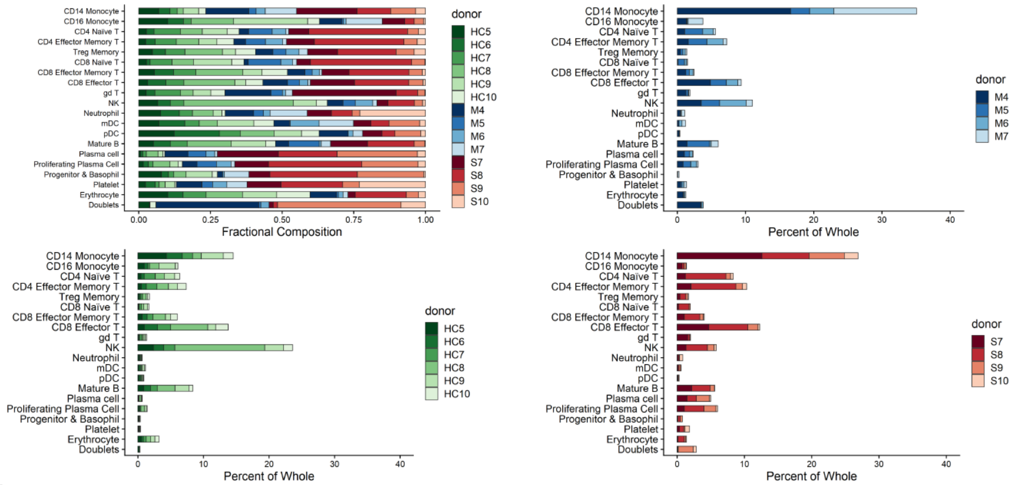
CD14 Monocyte
CD16 Monocyte
CD4 Naive T
CD4 Effector Memory T
Treg Memory
CD8 Naive T
CD8 Effector Memory T
CD8 Effector T
gd T
NK
Neutrophil
mDC
pDC
Mature B
Plasma cell
Proliferating Plasma Cell
Progenitor & Basophil
Platelet
Erythrocyte
Doublets
Fractional Composition
Percent of Whole
donor
HC5
HC6
HC7
HC8
HC9
HC10
M4
M5
M6
M7
S7
S8
S9
S10

O

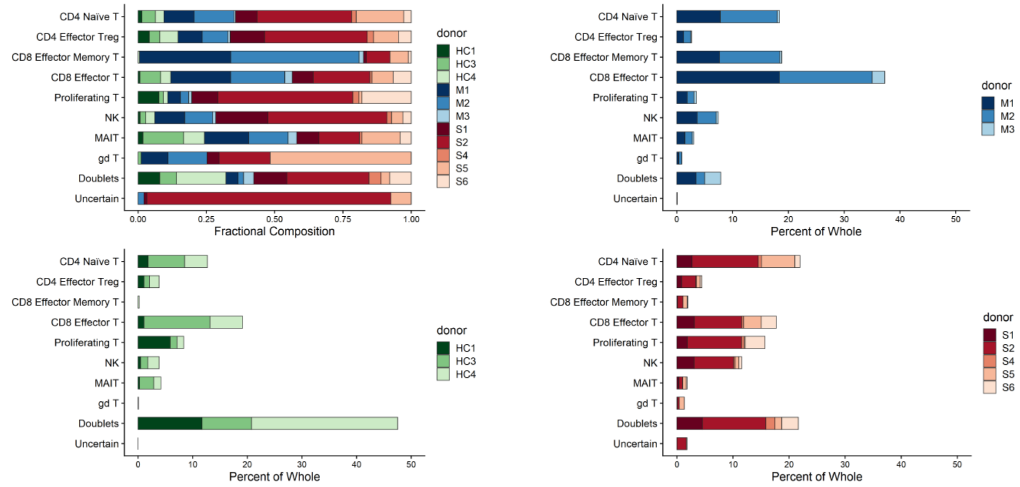
CD4 Naive T
CD4 Effector Treg
CD8 Effector Memory T
CD8 Effector T
Proliferating T
NK
MAIT
gd T
Doublets
Uncertain
Fractional Composition
Percent of Whole
donor
HC1
HC3
HC4
M1
M2
M3
S1
S2
S4
S5
S6

P

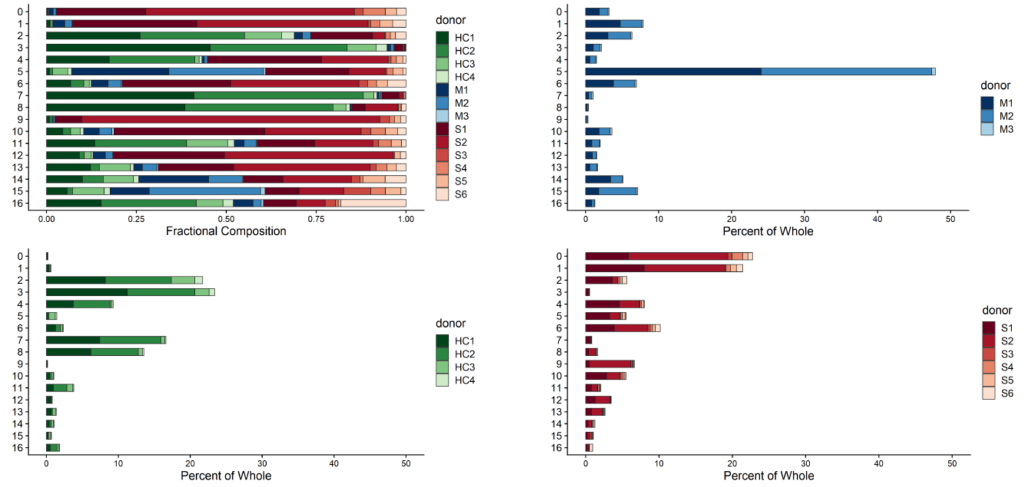
Fractional Composition
Percent of Whole
donor
HC1
HC2
HC3
HC4
M1
M2
M3
S1
S2
S3
S4
S5
S6

Q

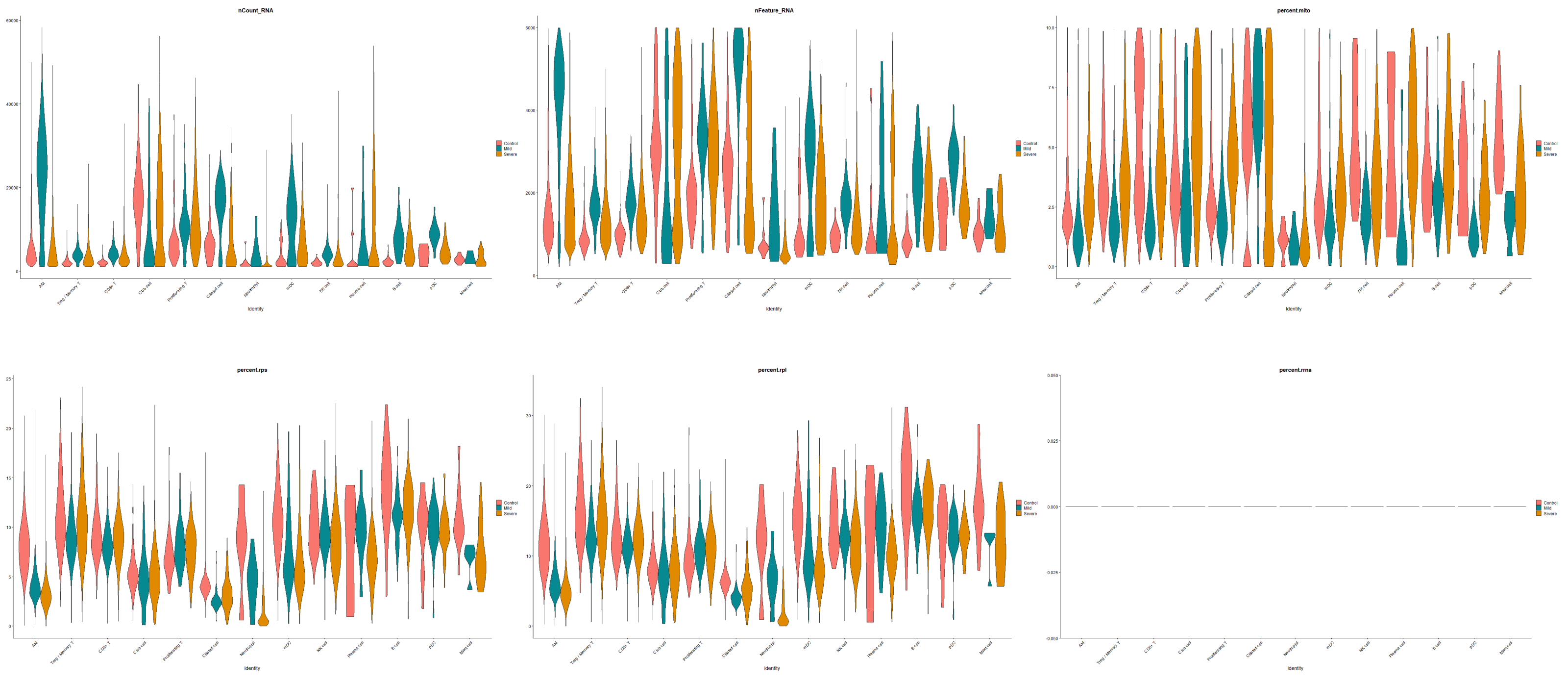
nCount_RNA
nFeature_RNA
percent.mito
percent.rps
percent.rpl
percent.rrna
Identity
Control
Mild
Severe

R

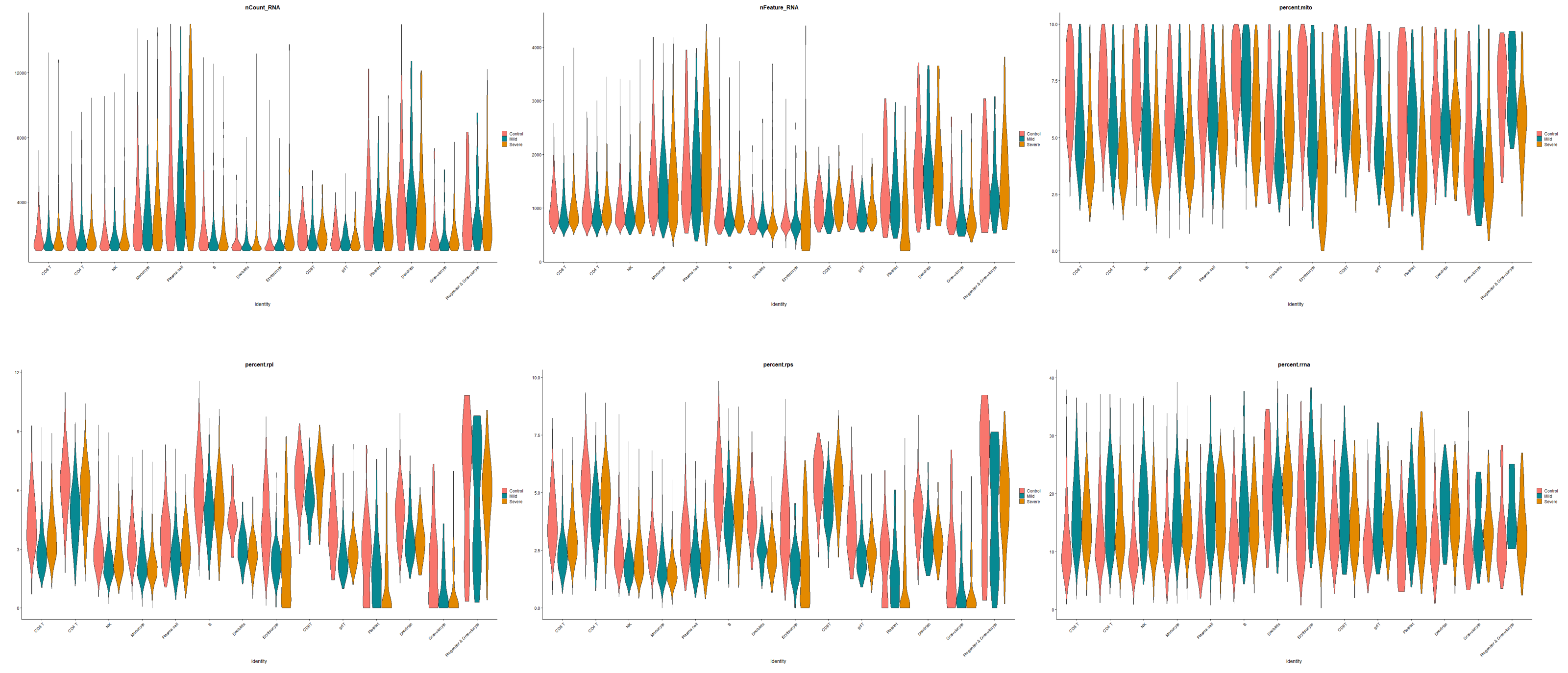
nCount_RNA
nFeature_RNA
percent.mito
percent.rpl
percent.rps
percent.rrna
Identity
Control
Mild
Severe

**Supplemental Figure S1.**

**A)** UMAP projection of integrated BALF data split by disease severity into groups of healthy control (*n*=4), moderate (*n*=3) and severe (*n*=6) donors.
**B)** Heatmap of differentially expressed marker genes used to determine BALF cluster identity.
**C)** Expression levels of canonical markers used for cell type annotation of the BALF clusters in Figure 1A.
**D)** UMAP projection of integrated PBMC data split by disease severity into groups of healthy control (*n*=6), non-ARDS (*n*=4) and ARDS (*n*=4) donors.
**E)** Heatmap of differentially expressed marker genes used to determine PBMC cluster identity.
**F)** Expression levels of canonical markers used for cell type annotation of the PBMC clusters in Figure 1B.
**G)** UMAP projection of NK and T cell populations following iterative data integration, split by disease severity into groups consisting of healthy controls (*n*=3), moderate (*n*=3), and severe (*n*=5) donors.
**H)** Heatmap of differentially expressed marker genes used to determine BALF NK and T cell cluster identity.
**I)** Expression levels of canonical markers used for cell type annotation of the clusters in Figure 1C.
**J)** UMAP projection of mononuclear phagocyte (MP) cell populations following iterative data integration, split by disease severity into groups consisting of healthy controls (*n*=4), moderate (*n*=3), and severe (*n*=6) donors. **K)** Heatmap of differentially expressed marker genes used to determine BALF MP cluster identity.
**L)** Expression levels of canonical markers used for detecting potential doublets following iterative data integration.
**M) (Top left)** Donor-specific fractional composition of BALF cell types in the total pool of healthy, moderate, and severe donors. **(Top right)** Donor-specific percentages of cell types recovered in the moderate donor pool. **(Bottom left)** Donor-specific percentages of cell types recovered in the healthy donor pool. **(Bottom right)** Donor-specific percentages of cell types recovered in the severe donor pool.
**N)** Donor-specific composition of PBMC cell types following the format of (M), but instead comparing healthy, non-ARDS, and ARDS donors.
**O)** Donor-specific composition of BALF NK and T cell types following the format of (M).
**P)** Donor-specific composition of BALF MPs following the format of (M).
**Q)** Quality control metrics for annotated BALF clusters. Top left to bottom right: Number of UMI counts, number of genes, percent mitochondrial transcripts, percent RPS ribosomal transcripts, percent RPL ribosomal transcripts, percent rRNA transcripts. rRNA transcripts were not identified in the BALF data.
**R)** Quality control metrics for annotated PBMC clusters. Top left to bottom right: Number of UMI counts, number of genes, percent mitochondrial transcripts, percent RPS ribosomal transcripts, percent RPL ribosomal transcripts, percent rRNA transcripts.

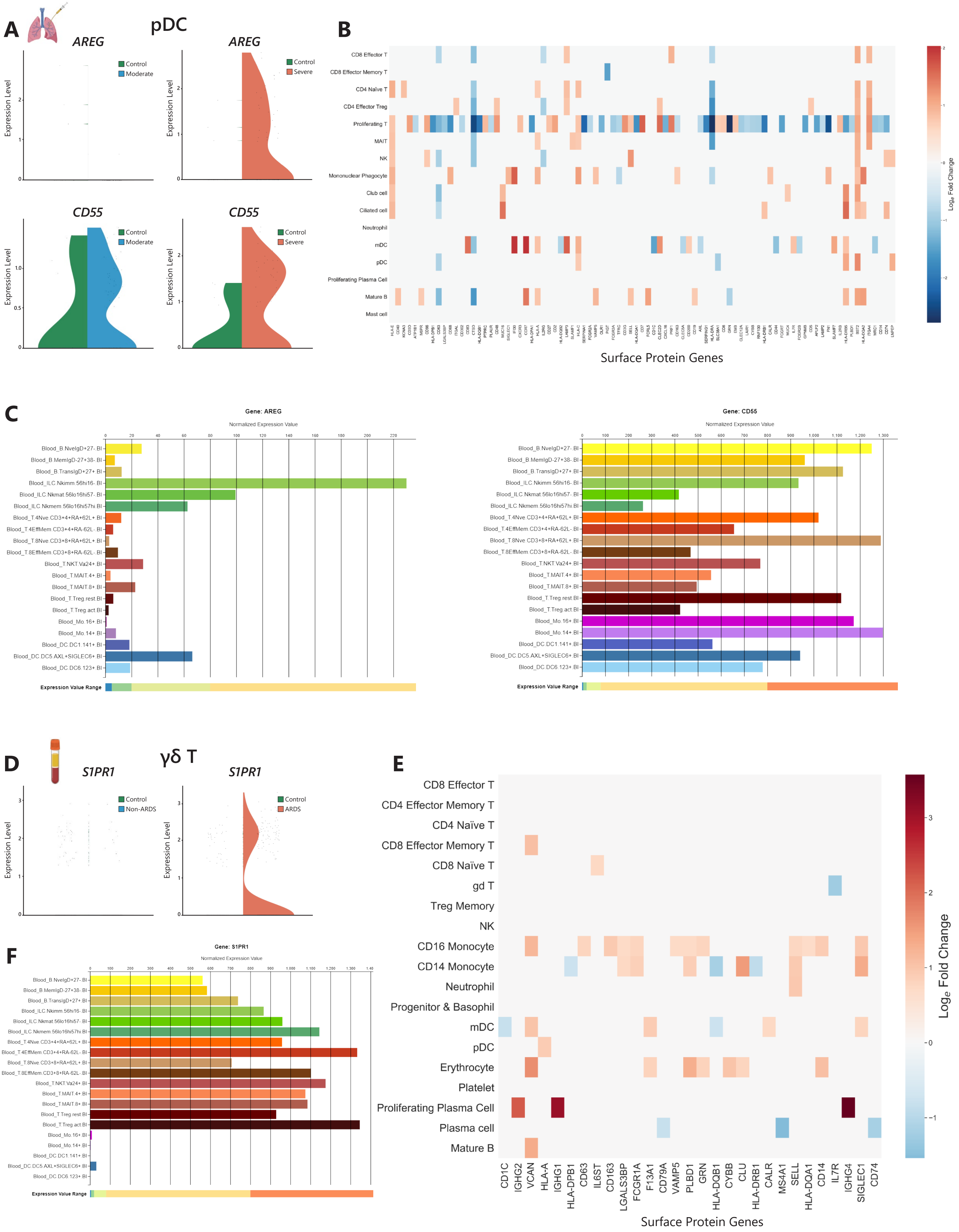
A
pDC
AREG
Control
Moderate
Expression Level
AREG
Control
Severe
CD55
Control
Moderate
CD55
Control
Severe
B
CD8 Effector T
CD8 Effector Memory T
CD4 Naïve T
CD4 Effector Treg
Proliferating T
MAIT
NK
Mononuclear Phagocyte
Club cell
Ciliated cell
Neutrophil
mDC
pDC
Proliferating Plasma Cell
Mature B
Mast cell
Surface Protein Genes
Loge Fold Change
C
Gene: AREG
Normalized Expression Value
Blood_B.NveIgD+27- Bl
Blood_B.MemIgD-27+38- Bl
Blood_B.TransIgD+27+ Bl
Blood_ILC.Nkimm.56hi16- Bl
Blood_ILC.Nkmat.56lo16hi57- Bl
Blood_ILC.Nkmem.56lo16hi57hi Bl
Blood_T.4Nve.CD3+4+RA+62L+ Bl
Blood_T.4EffMem.CD3+4+RA-62L- Bl
Blood_T.8Nve.CD3+8+RA+62L+ Bl
Blood_T.8EffMem.CD3+8+RA-62L- Bl
Blood_T.NKT.Va24+ Bl
Blood_T.MAIT.4+ Bl
Blood_T.MAIT.8+ Bl
Blood_T.Treg.rest.Bl
Blood_T.Treg.act.Bl
Blood_Mo.16+ Bl
Blood_Mo.14+ Bl
Blood_DC.DC1.141+ Bl
Blood_DC.DC5.AXL+SIGLEC6+ Bl
Blood_DC.DC6.123+ Bl
Expression Value Range
Gene: CD55
Normalized Expression Value
Expression Value Range
D
γδ T
S1PR1
Control
Non-ARDS
S1PR1
Control
ARDS
Expression Level
E
CD8 Effector T
CD4 Effector Memory T
CD4 Naïve T
CD8 Effector Memory T
CD8 Naïve T
gd T
Treg Memory
NK
CD16 Monocyte
CD14 Monocyte
Neutrophil
Progenitor & Basophil
mDC
pDC
Erythrocyte
Platelet
Proliferating Plasma Cell
Plasma cell
Mature B
CD1C
IGHG2
VCAN
HLA-A
IGHG1
HLA-DPB1
CD63
IL6ST
CD163
LGALS3BP
FCGR1A
F13A1
CD79A
VAMP5
PLBD1
GRN
HLA-DQB1
CYBB
CLU
HLA-DRB1
CALR
MS4A1
SELL
HLA-DQA1
CD14
IL7R
IGHG4
SIGLEC1
CD74
Surface Protein Genes
Loge Fold Change
F
Gene: S1PR1
Normalized Expression Value
Expression Value Range

G

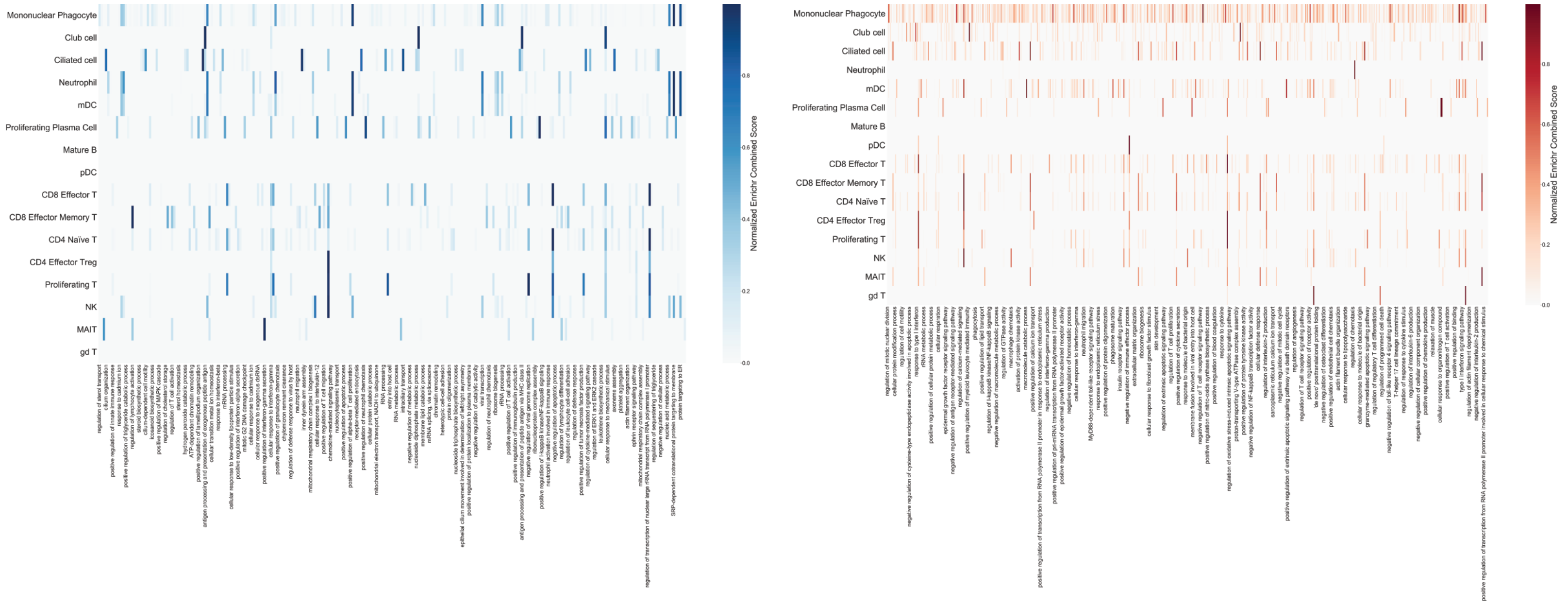
Mononuclear Phagocyte
Club cell
Ciliated cell
Neutrophil
mDC
Proliferating Plasma Cell
Mature B
pDC
CD8 Effector T
CD8 Effector Memory T
CD4 Naïve T
CD4 Effector Treg
Proliferating T
NK
MAIT
gd T
Normalized Enrichr Combined Score

H

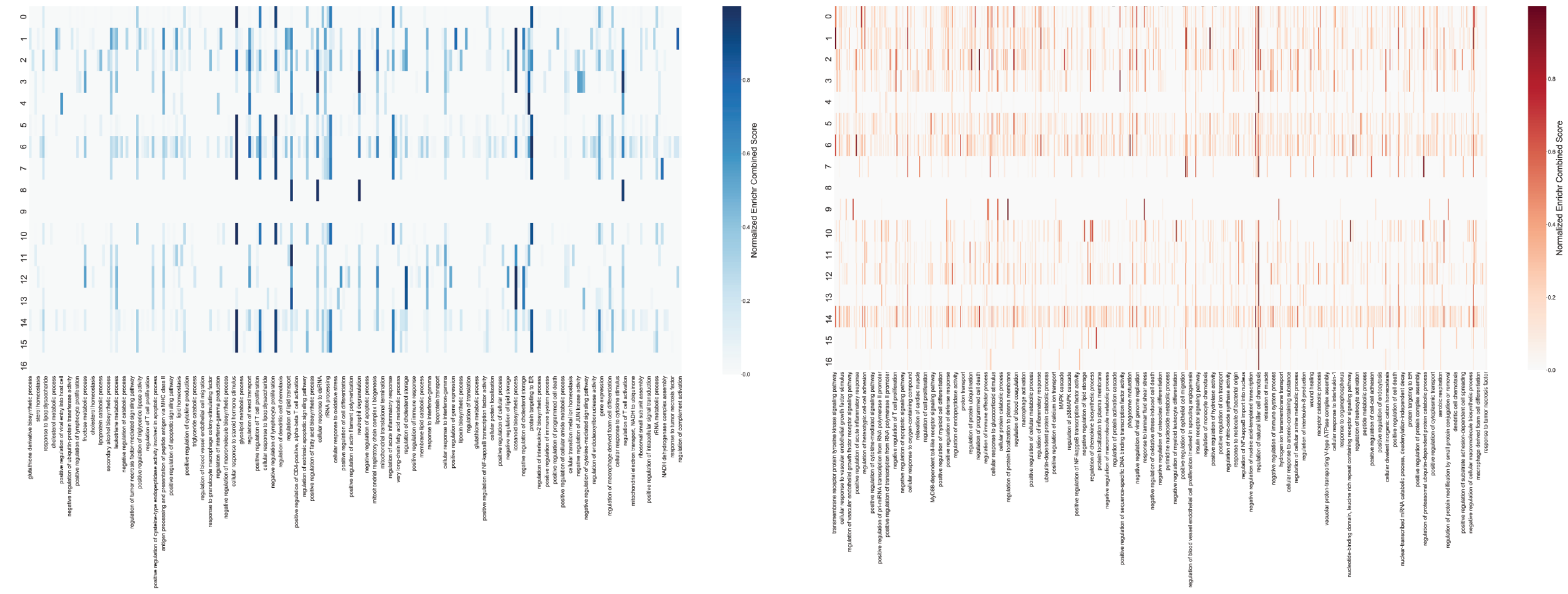
Normalized Enrichr Combined Score

I

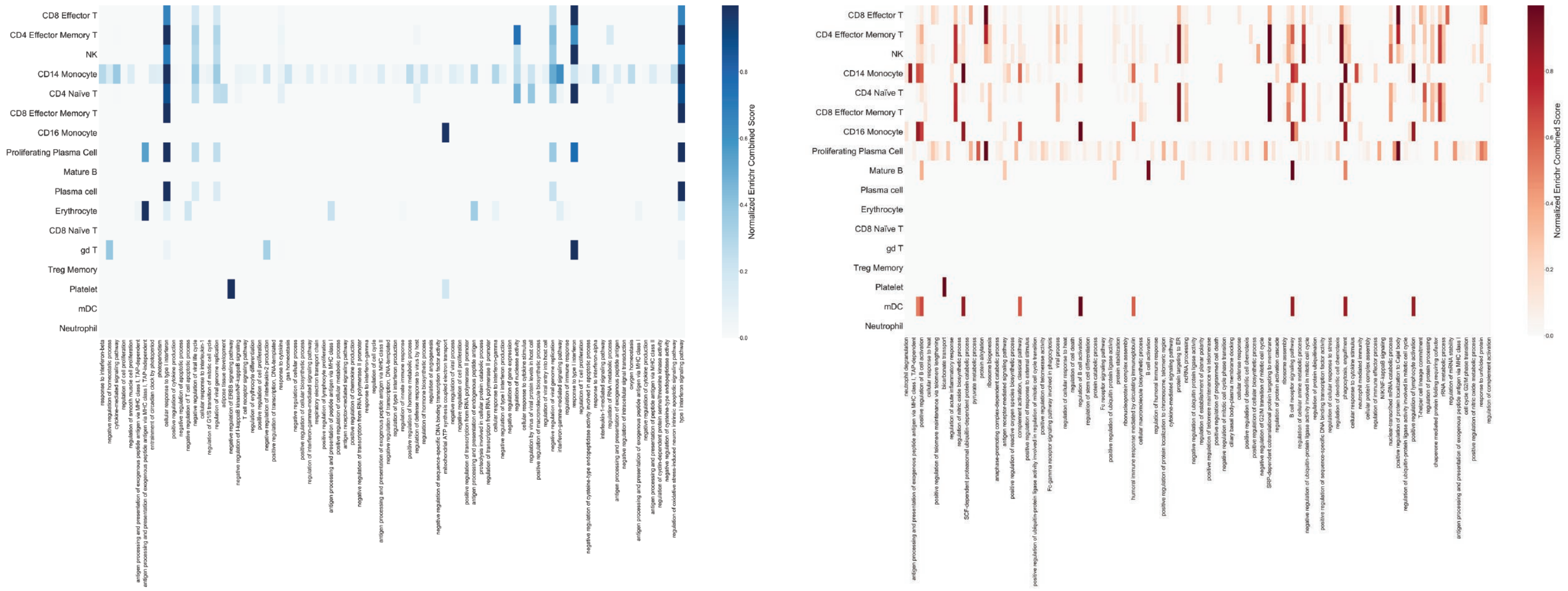
CD8 Effector T
CD4 Effector Memory T
NK
CD14 Monocyte
CD4 Naïve T
CD8 Effector Memory T
CD16 Monocyte
Proliferating Plasma Cell
Mature B
Plasma cell
Erythrocyte
CD8 Naïve T
gd T
Treg Memory
Platelet
mDC
Neutrophil
Normalized Enrichr Combined Score

**Supplemental Figure 2.**

**A)** Violin plots showing the differential expression of *AREG* and *CD55* in BALF pDCs when comparing control (green violins, *n*=3 for NK and T cells, *n*=4 for all other cell types) and moderate patients (blue violins, *n*=3) **(left)** and control and severe (red violins, *n*=5 for NK and T cells, *n*=6 for all others) patients **(right)**.

**B)** Heatmap displaying all cell surface markers found to be significantly ($|log_2FC|>1$ and $p<0.05$) differentially expressed in the BALF when comparing control and moderate patients.

**C)** Verification of *AREG* and *CD55* expression in blood pDCs through bulk RNA-seq data in the Immune Cell Atlas.

**D)** Violin plots showing the differential expression of *S1PR1* in γδ T cells of the PBMC donors when comparing control (*n*=6) and non-ARDS (*n*=4) patients **(left)** and control and ARDS (*n*=4) patients **(right)**.

**E)** Heatmap displaying all cell surface markers found to be significantly ($|log_2FC|>1$ and $p<0.05$) differentially expressed in PBMCs when comparing control and non-ARDS patients.

**F)** Verification of *S1PR1* expression in blood γδ T cells through bulk RNA-seq data in the Immune Cell Atlas.

**G)** Gene ontology (GO) annotation of differentially expressed genes according to biological functions for genes downregulated in severe vs. moderate disease in the BALF (including NK and T cell types).

**H)** GO annotation for genes upregulated in severe vs. moderate disease in the BALF.

**I)** GO annotation for genes downregulated in severe vs. moderate disease in the macrophage clusters from the BALF.

**J)** GO annotation for genes upregulated in severe vs. moderate disease in the macrophage clusters from the BALF.

**K)** GO annotation for genes downregulated in ARDS vs. non-ARDS in PBMCs.

**L)** GO annotation for genes upregulated in ARDS vs. non-ARDS in PBMCs.

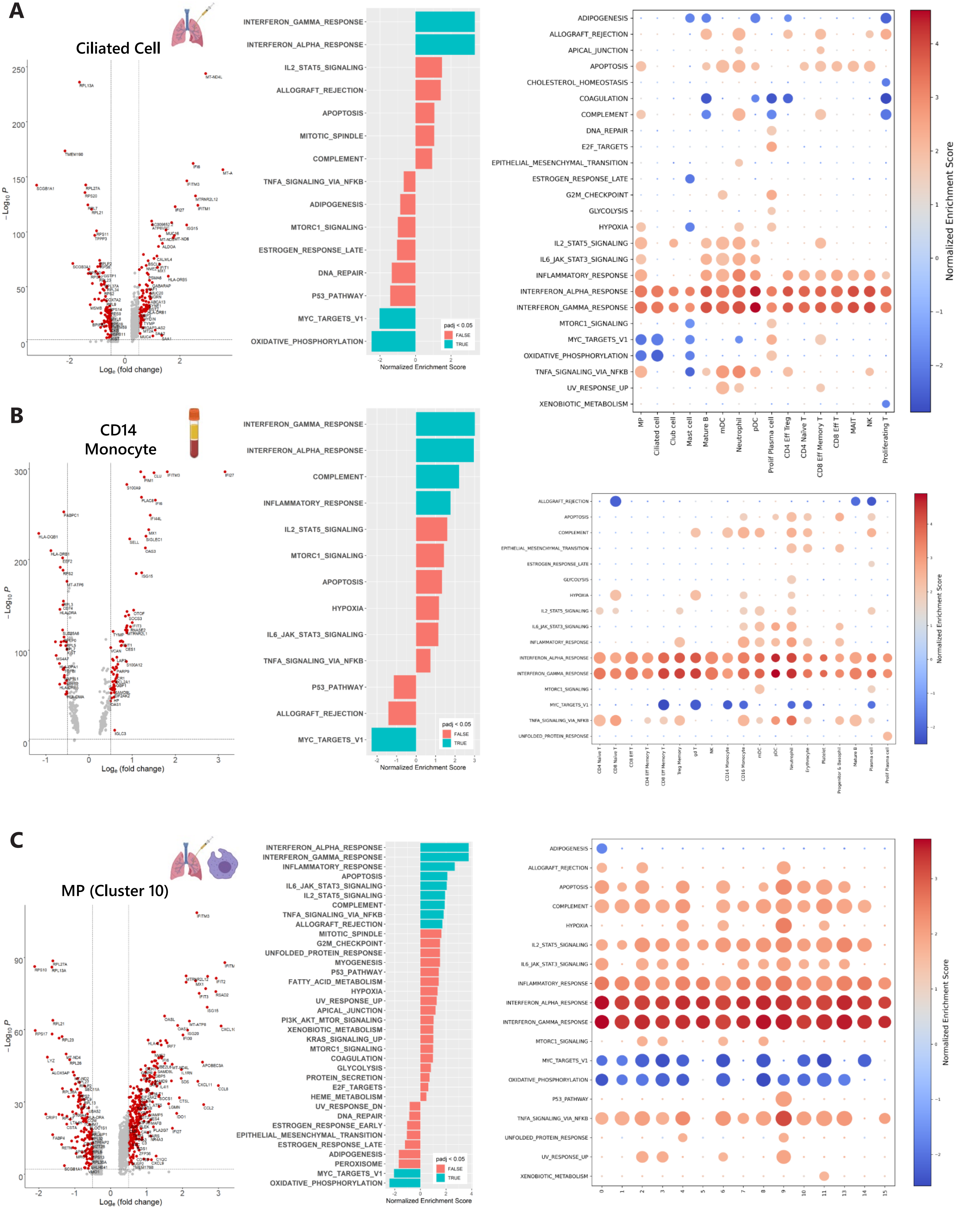
A
Ciliated Cell
−Log10 P
Loge (fold change)
INTERFERON_GAMMA_RESPONSE
INTERFERON_ALPHA_RESPONSE
IL2_STAT5_SIGNALING
ALLOGRAFT_REJECTION
APOPTOSIS
MITOTIC_SPINDLE
COMPLEMENT
TNFA_SIGNALING_VIA_NFKB
ADIPOGENESIS
MTORC1_SIGNALING
ESTROGEN_RESPONSE_LATE
DNA_REPAIR
P53_PATHWAY
MYC_TARGETS_V1
OXIDATIVE_PHOSPHORYLATION
padj < 0.05
FALSE
TRUE
Normalized Enrichment Score
ADIPOGENESIS
ALLOGRAFT_REJECTION
APICAL_JUNCTION
APOPTOSIS
CHOLESTEROL_HOMEOSTASIS
COAGULATION
COMPLEMENT
DNA_REPAIR
E2F_TARGETS
EPITHELIAL_MESENCHYMAL_TRANSITION
ESTROGEN_RESPONSE_LATE
G2M_CHECKPOINT
GLYCOLYSIS
HYPOXIA
IL2_STAT5_SIGNALING
IL6_JAK_STAT3_SIGNALING
INFLAMMATORY_RESPONSE
INTERFERON_ALPHA_RESPONSE
INTERFERON_GAMMA_RESPONSE
MTORC1_SIGNALING
MYC_TARGETS_V1
OXIDATIVE_PHOSPHORYLATION
TNFA_SIGNALING_VIA_NFKB
UV_RESPONSE_UP
XENOBIOTIC_METABOLISM
MP
Ciliated cell
Club cell
Mast cell
Mature B
mDC
Neutrophil
pDC
Prolif Plasma cell
CD4 Eff Treg
CD4 Naive T
CD8 Eff Memory T
CD8 Eff T
MAIT
NK
Proliferating T
Normalized Enrichment Score
B
CD14
Monocyte
−Log10 P
Loge (fold change)
INTERFERON_GAMMA_RESPONSE
INTERFERON_ALPHA_RESPONSE
COMPLEMENT
INFLAMMATORY_RESPONSE
IL2_STAT5_SIGNALING
MTORC1_SIGNALING
APOPTOSIS
HYPOXIA
IL6_JAK_STAT3_SIGNALING
TNFA_SIGNALING_VIA_NFKB
P53_PATHWAY
ALLOGRAFT_REJECTION
MYC_TARGETS_V1
padj < 0.05
FALSE
TRUE
Normalized Enrichment Score
ALLOGRAFT_REJECTION
APOPTOSIS
COMPLEMENT
EPITHELIAL_MESENCHYMAL_TRANSITION
ESTROGEN_RESPONSE_LATE
GLYCOLYSIS
HYPOXIA
IL2_STAT5_SIGNALING
IL6_JAK_STAT3_SIGNALING
INFLAMMATORY_RESPONSE
INTERFERON_ALPHA_RESPONSE
INTERFERON_GAMMA_RESPONSE
MTORC1_SIGNALING
MYC_TARGETS_V1
TNFA_SIGNALING_VIA_NFKB
UNFOLDED_PROTEIN_RESPONSE
CD4 Naive T
CD8 Naive T
CD8 Eff T
CD4 Eff Memory T
CD8 Eff Memory T
Treg Memory
gd T
NK
CD14 Monocyte
CD16 Monocyte
mDC
pDC
Neutrophil
Erythrocyte
Platelet
Progenitor & Basophil
Mature B
Plasma cell
Prolif Plasma cell
Normalized Enrichment Score
C
MP (Cluster 10)
−Log10 P
Loge (fold change)
INTERFERON_ALPHA_RESPONSE
INTERFERON_GAMMA_RESPONSE
INFLAMMATORY_RESPONSE
APOPTOSIS
IL6_JAK_STAT3_SIGNALING
IL2_STAT5_SIGNALING
COMPLEMENT
TNFA_SIGNALING_VIA_NFKB
ALLOGRAFT_REJECTION
MITOTIC_SPINDLE
G2M_CHECKPOINT
UNFOLDED_PROTEIN_RESPONSE
MYOGENESIS
P53_PATHWAY
FATTY_ACID_METABOLISM
HYPOXIA
UV_RESPONSE_UP
APICAL_JUNCTION
PI3K_AKT_MTOR_SIGNALING
XENOBIOTIC_METABOLISM
KRAS_SIGNALING_UP
MTORC1_SIGNALING
COAGULATION
GLYCOLYSIS
PROTEIN_SECRETION
E2F_TARGETS
HEME_METABOLISM
UV_RESPONSE_DN
DNA_REPAIR
ESTROGEN_RESPONSE_EARLY
EPITHELIAL_MESENCHYMAL_TRANSITION
ESTROGEN_RESPONSE_LATE
ADIPOGENESIS
PEROXISOME
MYC_TARGETS_V1
OXIDATIVE_PHOSPHORYLATION
padj < 0.05
FALSE
TRUE
Normalized Enrichment Score
ADIPOGENESIS
ALLOGRAFT_REJECTION
APOPTOSIS
COMPLEMENT
HYPOXIA
IL2_STAT5_SIGNALING
IL6_JAK_STAT3_SIGNALING
INFLAMMATORY_RESPONSE
INTERFERON_ALPHA_RESPONSE
INTERFERON_GAMMA_RESPONSE
MTORC1_SIGNALING
MYC_TARGETS_V1
OXIDATIVE_PHOSPHORYLATION
P53_PATHWAY
TNFA_SIGNALING_VIA_NFKB
UNFOLDED_PROTEIN_RESPONSE
UV_RESPONSE_UP
XENOBIOTIC_METABOLISM
Normalized Enrichment Score

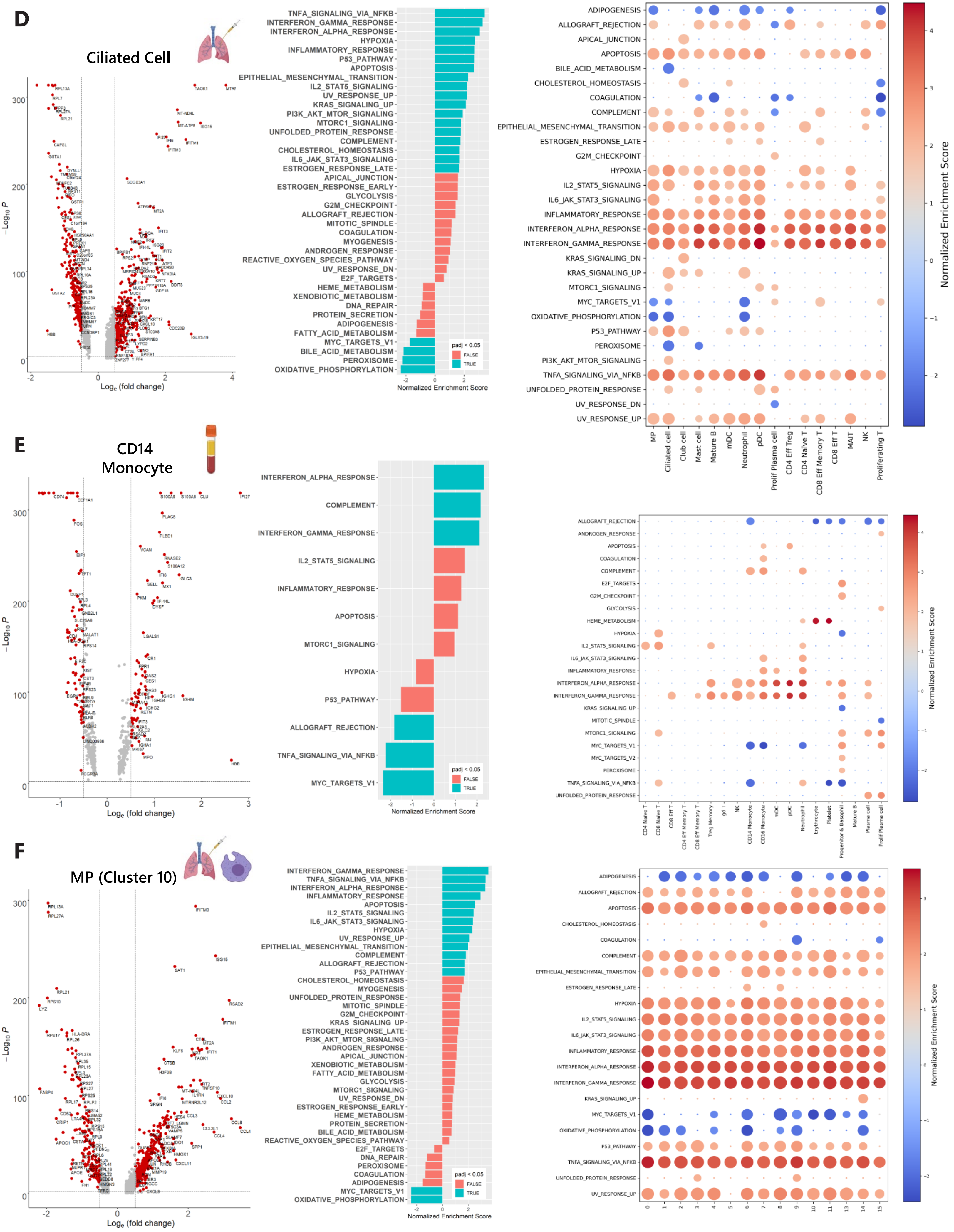

D
Ciliated Cell
−Log10 P
Loge (fold change)
TNFA_SIGNALING_VIA_NFKB
INTERFERON_GAMMA_RESPONSE
INTERFERON_ALPHA_RESPONSE
HYPOXIA
INFLAMMATORY_RESPONSE
P53_PATHWAY
APOPTOSIS
EPITHELIAL_MESENCHYMAL_TRANSITION
IL2_STAT5_SIGNALING
UV_RESPONSE_UP
KRAS_SIGNALING_UP
PI3K_AKT_MTOR_SIGNALING
MTORC1_SIGNALING
UNFOLDED_PROTEIN_RESPONSE
COMPLEMENT
CHOLESTEROL_HOMEOSTASIS
IL6_JAK_STAT3_SIGNALING
ESTROGEN_RESPONSE_LATE
APICAL_JUNCTION
ESTROGEN_RESPONSE_EARLY
GLYCOLYSIS
G2M_CHECKPOINT
ALLOGRAFT_REJECTION
MITOTIC_SPINDLE
COAGULATION
MYOGENESIS
ANDROGEN_RESPONSE
REACTIVE_OXYGEN_SPECIES_PATHWAY
UV_RESPONSE_DN
E2F_TARGETS
HEME_METABOLISM
XENOBIOTIC_METABOLISM
DNA_REPAIR
PROTEIN_SECRETION
ADIPOGENESIS
FATTY_ACID_METABOLISM
MYC_TARGETS_V1
BILE_ACID_METABOLISM
PEROXISOME
OXIDATIVE_PHOSPHORYLATION
padj < 0.05
FALSE
TRUE
Normalized Enrichment Score
MP
Ciliated cell
Club cell
Mast cell
Mature B
mDC
Neutrophil
pDC
Prolif Plasma cell
CD4 Eff Treg
CD4 Naive T
CD8 Eff Memory T
CD8 Eff T
MAIT
NK
Proliferating T
Normalized Enrichment Score
E
CD14 Monocyte
INTERFERON_ALPHA_RESPONSE
COMPLEMENT
INTERFERON_GAMMA_RESPONSE
IL2_STAT5_SIGNALING
INFLAMMATORY_RESPONSE
APOPTOSIS
MTORC1_SIGNALING
HYPOXIA
P53_PATHWAY
ALLOGRAFT_REJECTION
TNFA_SIGNALING_VIA_NFKB
MYC_TARGETS_V1
padj < 0.05
FALSE
TRUE
Normalized Enrichment Score
CD4 Naive T
CD8 Naive T
CD8 Eff T
CD4 Eff Memory T
CD8 Eff Memory T
Treg Memory
gd T
NK
CD14 Monocyte
CD16 Monocyte
mDC
pDC
Neutrophil
Erythrocyte
Platelet
Progenitor & Basophil
Mature B
Plasma cell
Prolif Plasma cell
Normalized Enrichment Score
F
MP (Cluster 10)
−Log10 P
Loge (fold change)
INTERFERON_GAMMA_RESPONSE
TNFA_SIGNALING_VIA_NFKB
INTERFERON_ALPHA_RESPONSE
INFLAMMATORY_RESPONSE
APOPTOSIS
IL2_STAT5_SIGNALING
IL6_JAK_STAT3_SIGNALING
HYPOXIA
UV_RESPONSE_UP
EPITHELIAL_MESENCHYMAL_TRANSITION
COMPLEMENT
ALLOGRAFT_REJECTION
P53_PATHWAY
CHOLESTEROL_HOMEOSTASIS
MYOGENESIS
UNFOLDED_PROTEIN_RESPONSE
MITOTIC_SPINDLE
G2M_CHECKPOINT
KRAS_SIGNALING_UP
ESTROGEN_RESPONSE_LATE
PI3K_AKT_MTOR_SIGNALING
ANDROGEN_RESPONSE
APICAL_JUNCTION
XENOBIOTIC_METABOLISM
FATTY_ACID_METABOLISM
GLYCOLYSIS
MTORC1_SIGNALING
UV_RESPONSE_DN
ESTROGEN_RESPONSE_EARLY
HEME_METABOLISM
PROTEIN_SECRETION
BILE_ACID_METABOLISM
REACTIVE_OXYGEN_SPECIES_PATHWAY
E2F_TARGETS
DNA_REPAIR
PEROXISOME
COAGULATION
ADIPOGENESIS
MYC_TARGETS_V1
OXIDATIVE_PHOSPHORYLATION
padj < 0.05
FALSE
TRUE
Normalized Enrichment Score
Normalized Enrichment Score

G

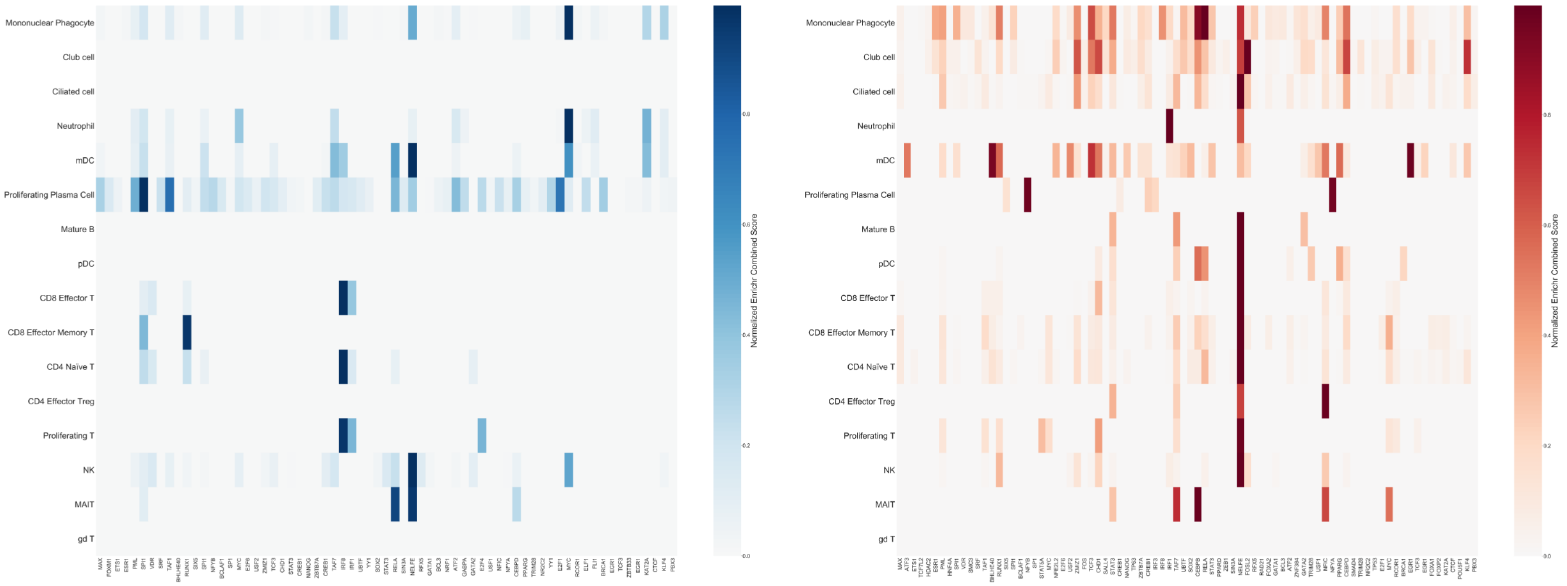
Mononuclear Phagocyte
Club cell
Ciliated cell
Neutrophil
mDC
Proliferating Plasma Cell
Mature B
pDC
CD8 Effector T
CD8 Effector Memory T
CD4 Naïve T
CD4 Effector Treg
Proliferating T
NK
MAIT
gd T
Normalized Enrichr Combined Score

H

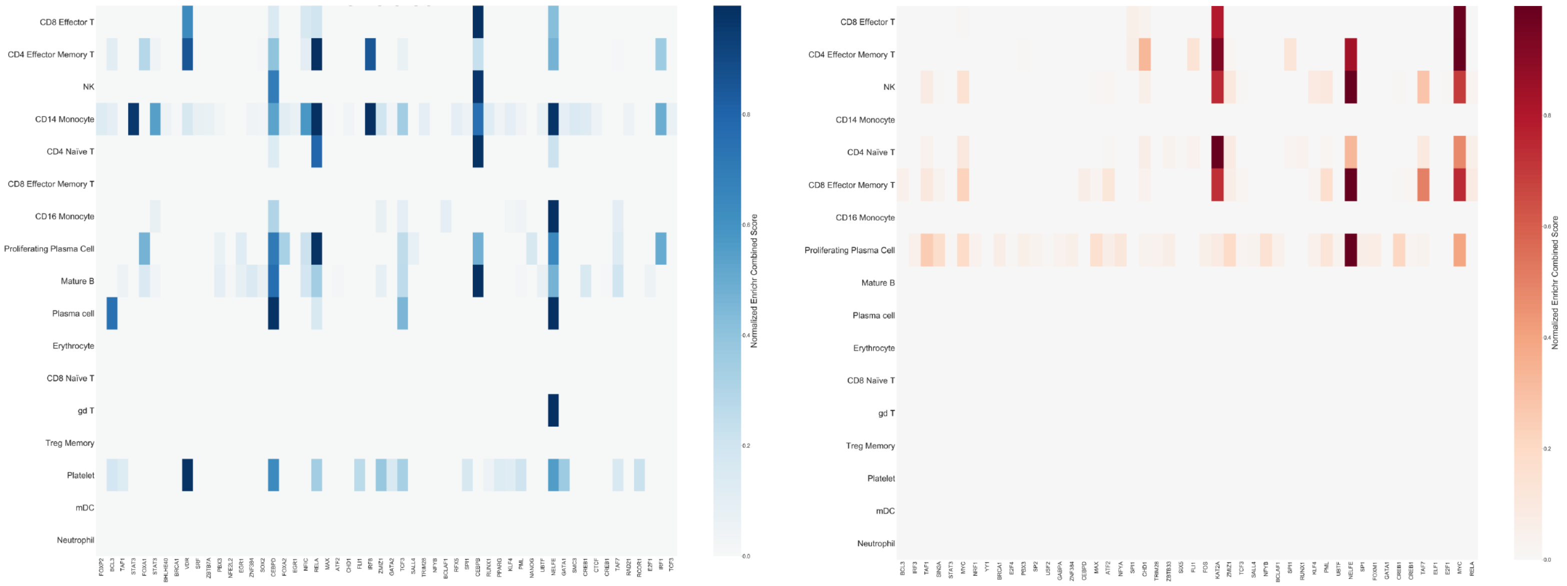
CD8 Effector T
CD4 Effector Memory T
NK
CD14 Monocyte
CD4 Naïve T
CD8 Effector Memory T
CD16 Monocyte
Proliferating Plasma Cell
Mature B
Plasma cell
Erythrocyte
CD8 Naïve T
gd T
Treg Memory
Platelet
mDC
Neutrophil
Normalized Enrichr Combined Score

I

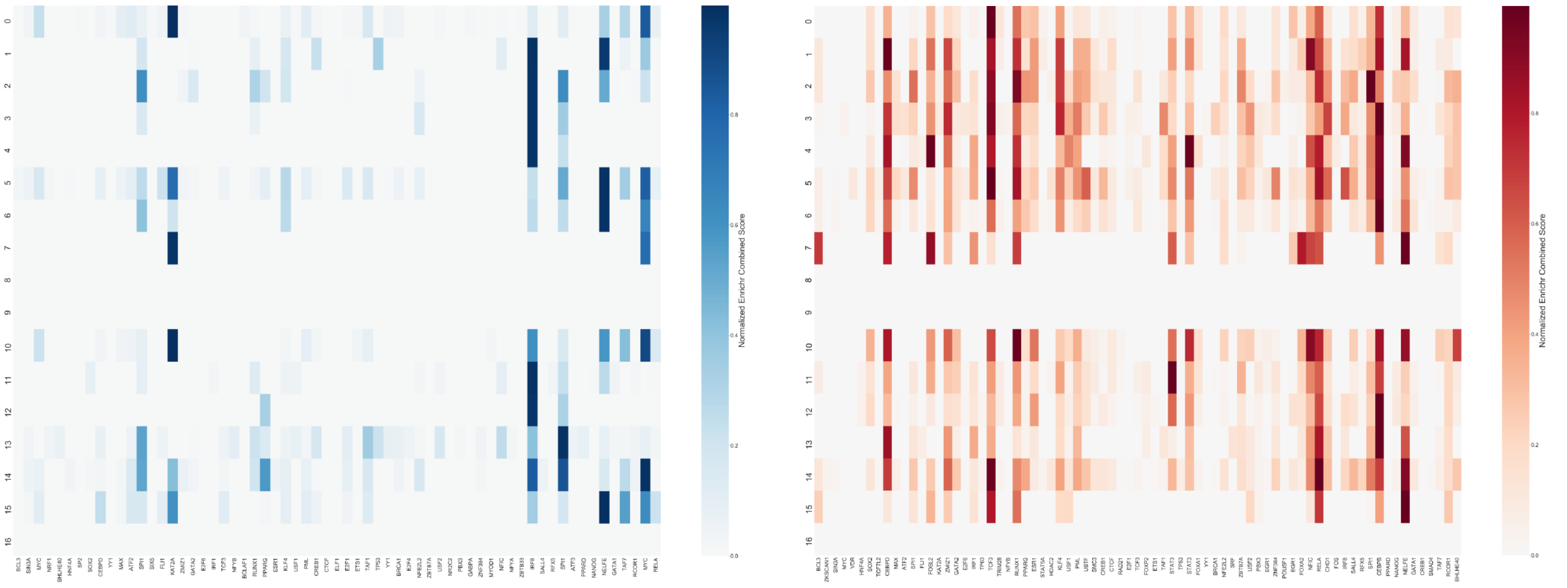
Normalized Enrichr Combined Score

**Supplemental Figure S3.**

**A) (Left)** Volcano plot showing significantly differentially expressed genes (red dots) in ciliated cells from BALF when comparing moderate and control patients. **(Middle)** Gene set enrichment analysis (GSEA) of hallmark gene sets in the ciliated cells of the lung. Bars colored blue are significantly enriched or depleted (adjusted $p<0.05$) while bars colored red do not meet the significance threshold. **(Right)** Hallmark gene sets indicate enrichment of molecular pathways for all cell types found in the BALF. Normalized enrichment score is shown by a blue-red color scale and dot size is proportional to $-\log_{10}(p)$, with the smallest dot size indicating non-significant adjusted p-values ($p<0.05$). MP=MPs.

**B) (Left)** Volcano plot showing significantly differentially expressed genes (red dots) in CD14+ monocytes from blood when comparing non-ARDS COVID-19 patients and control subjects. **(Middle)** Gene set enrichment analysis (GSEA) of hallmark gene sets in CD14+ monocytes. **(Right)** Hallmark gene sets indicate significant pathway enrichment in ARDS for all PBMC cell types found in the blood.

**C) (Left)** Volcano plot showing significantly differentially expressed genes (red dots) in the one of 16 mononuclear phagocyte (MP) populations (cluster 10) from the BALF when comparing moderate and control patients. **(Middle)** Gene set enrichment analysis (GSEA) of hallmark gene sets in cluster 10 of BALF MPs. **(Right)** Hallmark gene sets indicate enrichment of molecular pathways for all MP clusters found in the BALF. Clusters 12 and 16 represent suspected doublets.

**D)** Comparison between BALF responses in severe patients and healthy controls following the format of (A).

**E)** Comparison between PBMC responses in ARDS patients and healthy controls following the format of (B).

**F)** Comparison between BALF MP responses in severe patients and healthy controls following the format of (C).

**G) (Left)** Heatmap displaying transcription factors found to be downregulated in the BALF of severe patients when compared to moderate patients**. (Right)** Heatmap displaying transcription factors found to be upregulated in the BALF of severe patients when compared to moderate patients.

**H) (Left)** Heatmap displaying transcription factors found to be downregulated in PBMCs of ARDS patients when compared to non-ARDS patients. **(Right)** Heatmap displaying transcription factors found to be upregulated in PBMCs of ARDS patients when compared to non-ARDS patients.

**I) (Left)** Heatmap displaying transcription factors found to be downregulated in the MPs in the BALF of severe patients when compared to moderate patients. **(Right)** Heatmap displaying transcription factors found to be upregulated in the MPs in the BALF of severe patients when compared to moderate patients

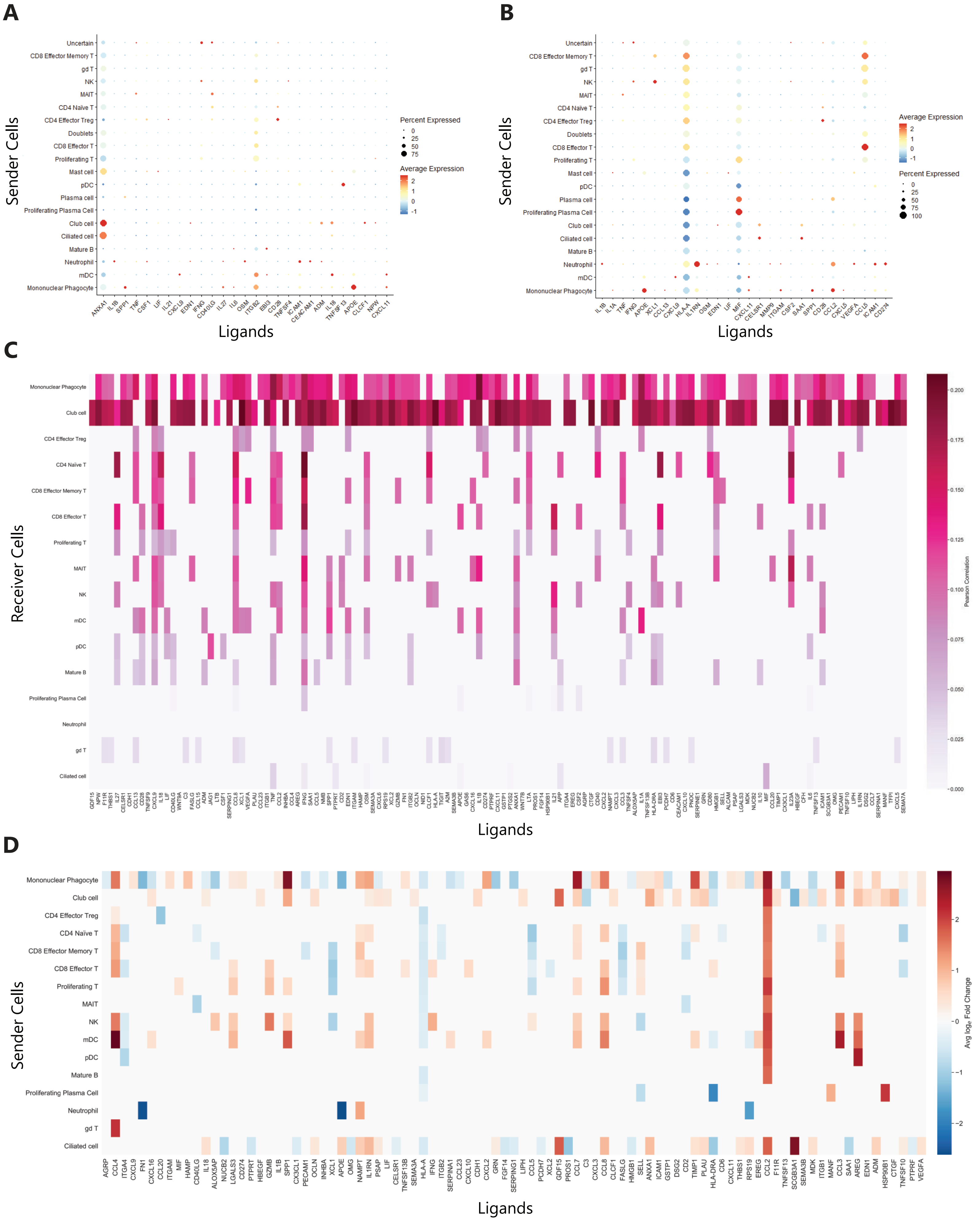
A
Sender Cells
Ligands
Percent Expressed
0
25
50
75
Average Expression
2
1
0
-1
Uncertain
CD8 Effector Memory T
gd T
NK
MAIT
CD4 Naïve T
CD4 Effector Treg
Doublets
CD8 Effector T
Proliferating T
Mast cell
pDC
Plasma cell
Proliferating Plasma Cell
Club cell
Ciliated cell
Mature B
Neutrophil
mDC
Mononuclear Phagocyte
B
Sender Cells
Ligands
Average Expression
2
1
0
-1
Percent Expressed
0
25
50
75
100
C
Receiver Cells
Ligands
Pearson Correlation
0.200
0.175
0.150
0.125
0.100
0.075
0.050
0.025
0.000
Mononuclear Phagocyte
Club cell
CD4 Effector Treg
CD4 Naïve T
CD8 Effector Memory T
CD8 Effector T
Proliferating T
MAIT
NK
mDC
pDC
Mature B
Proliferating Plasma Cell
Neutrophil
gd T
Ciliated cell
D
Sender Cells
Ligands
Avg loge Fold Change
2
1
0
−1
−2

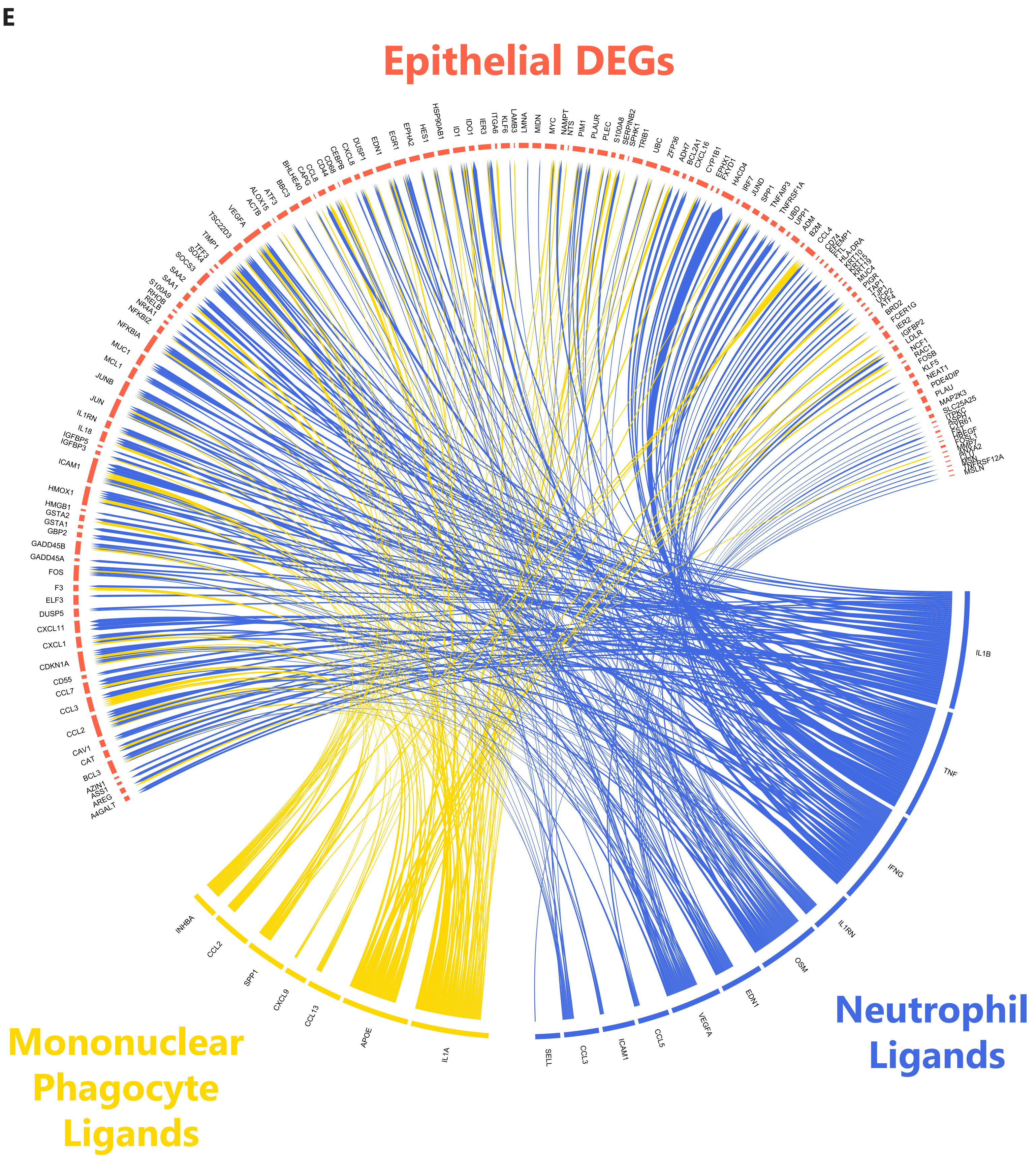

E
Epithelial DEGs
Neutrophil Ligands
Mononuclear Phagocyte Ligands
IL1B
TNF
IFNG
IL1RN
OSM
EDN1
VEGFA
CCL5
ICAM1
CCL3
SELL
IL1A
APOE
CCL13
CXCL9
SPP1
CCL2
INHBA
A4GALT
AREG
ASS1
AZIN1
BCL3
CAT
CAV1
CCL2
CCL3
CCL7
CD55
CDKN1A
CXCL1
CXCL11
DUSP5
ELF3
F3
FOS
GADD45A
GADD45B
GBP2
GSTA1
GSTA2
HMGB1
HMOX1
ICAM1
IGFBP3
IGFBP5
IL18
IL1RN
JUN
JUNB
MCL1
MUC1
NFKBIA
NFKBIZ
NR4A1
RELB
RHOB
S100A9
SAA1
SAA2
SOCS3
SOX4
TFF3
TIMP1
TSC22D3
VEGFA
ACTB
ALOX15
ATF3
BBC3
BHLHE40
CAPG
CCL8
CD44
CD68
CEBPB
CXCL8
DUSP1
EDN1
EGR1
EPHA2
HES1
HSP90AB1
ID1
IDO1
IER3
ITGA6
KLF6
LAMB3
LMNA
MIDN
MYC
NAMPT
NTS
PIM1
PLAUR
PLEC
S100A8
SERPINB2
SPHK1
TRIB1
UBC
ZFP36
ADH7
BCL2A1
CXCL16
CYP1B1
EPHX1
FXYD1
HACD4
IRF7
JUND
SPP1
TNFAIP3
TNFRSF1A
UBD
UPP1
ADM
B2M
CCL4
CD74
HLA-DRA
KRT19
MUC4
PIGR
ATF4
BRD2
FCER1G
IER2
IGFBP2
LDLR
NCF1
RAC1
FOSB
KLF5
NEAT1
PDE4DIP
PLAU
MAP2K3
SLC25A25
MSLN

**Supplemental Figure S4.**

**A)** Dotplot showing BALF "sender" cells expressing ligands implicated in the differential expression observed in pDCs. Dot size indicates the percent of the population that is expressing the ligand and the red-blue color scale indicates the average normalized expression level.
**B)** Dotplot showing BALF "sender" cells expressing ligands implicated in the differential expression observed in epithelial cells. Dot size indicates the percent of the population that is expressing the ligand and the red-blue color scale indicates the average normalized expression level.
**C)** Full list of NicheNet-identified ligands potentially driving differential expression in "receiver" cells of the BALF. The pink color scale indicates Pearson correlation, a measure of a ligand's ability to induce the differential expression observed in the corresponding "receiver" cell.
**D)** Differential expression of NicheNet-identified ligands between severe and moderate disease in all potential "sender" cells of the BALF. Ligands that are significantly differentially expressed in severe disease with adjusted $p<0.05$ are colored by their average natural log fold change on a blue-red color scale.
**E)** Expansion of the circos plot found in (C) showing ligand-target relationships between target DEGs in epithelial cells (red) and ligands expressed by mononuclear phagocytes (yellow) and neutrophils (blue).
